# Homeostatic regulation of neuronal activity: implications and feasibility of compartment-specific feedback

**DOI:** 10.64898/2025.12.10.693436

**Authors:** Ankit Roy, Steven A. Prescott

**Affiliations:** Neurosciences and Mental Health, The Hospital for Sick Children, Toronto, ON, Canada; Department of Physiology, University of Toronto, Toronto, ON, Canada; Institute of Biomedical Engineering, University of Toronto, Toronto, ON, Canada; Department of Physiology and Pharmacology, University of Calgary, Calgary, AB, Canada; Hotchkiss Brain Institute, University of Calgary, Calgary, AB, Canada

## Abstract

Neurons regulate their average firing rate by adjusting their synaptic strength and intrinsic excitability. Intracellular calcium is implicated as an error signal for both types of homeostatic regulation. Past studies have focused on global calcium but two properties cannot be independently regulated by one error signal. Here, we computationally tested the implications of local vs global feedback and the feasibility of local (compartment-specific) error signals based on spatially segregated calcium changes. Simulations in a simple two-compartment model confirmed that a perturbation applied to one compartment induces local compensation only if feedback is compartment-specific. Simulations in a biophysically detailed multicompartment model with realistic calcium handling confirmed that dendritic and somatic calcium signals remain relatively segregated and can, therefore, encode separate error signals. Strong perturbations (as often tested experimentally) triggered widespread compensation because local compensation was overwhelmed. Non-local compensation also occurred when the spatial segregation of calcium signals was weakened. Our results demonstrate the plausibility of compartment-specific feedback using calcium-based error signals. Furthermore, whereas local homeostatic regulation nullifies local perturbations through compensation within the affected compartment, non-local regulation causes widespread compensatory changes that, while restoring the neuron’s overall input-output relationship, distorts the input-output relationship of individual compartments, with potentially important consequences.

## INTRODUCTION

Neurons use action potentials, or spikes, to encode and transmit information. Firing rate varies on short timescales due to changing input but, on longer timescales, neurons maintain their average firing rate near a target value by adjusting ion channel densities to offset chronic changes in input or other factors. Compensatory changes in synaptic strength or intrinsic excitability are referred to as synaptic scaling and excitability regulation, respectively^1,2^. Such regulation occurs in response to perturbations such as chronic activity blockade^3^, long-term synaptic changes^4,5^ and chronic somatic stimulation ^6^, enabling neurons to continue operating within their dynamic range^7^.

Intracellular calcium is affected by synaptic input and neuronal spiking, and can thus encode activity levels. Furthermore, calcium is involved in regulating diverse neuronal processes^8^. Sustained calcium changes activate calmodulin (CaM), a ubiquitous calcium sensor^9^. The Ca^2+^/CaM complex can bind with diverse protein kinases called CaMKs, including CaMKII which regulates synaptic strength and intrinsic excitability via transcriptional changes^10^ and CaMKIII which acts by modulating translation^9^. Blocking calcium influx through voltage-gated calcium channels (VGCCs) induces synaptic upscaling^11^. In brief, chronic changes in activity trigger compensatory adjustments via deviations in “resting” calcium levels.

Calcium-mediated feedback has been used in several computational models to regulate average firing rate via activity-dependent adjustment of channel densities^12–17^. These studies implicitly focussed on global calcium changes since neurons were homogeneous single-compartmental models. However, evidence suggests activity-dependent changes can occur independently at different locations within the same neuron^6,18–20^. Moreover, division of a complex system into modules with independent feedback controllers increases flexibility and stability of the system to multiple inputs^21–23^. Thus, if a neuron receives chronically increased or decreased synaptic input, or if its spiking output is otherwise chronically altered, offsetting the specific effects of a perturbation with targeted compensation is likely preferable to broader, less specific compensation. Independently regulating synaptic strength and excitability by adjusting dendritic and somatic ion channel densities, respectively, would enable a neuron to nullify a perturbation locally and avoid broader changes that risk introducing collateral disruptions. One requirement for local regulation is compartment-specific feedback, i.e. a local error signal. The calcium rise caused by synaptic activation is known to remain restricted to areas near the synapse, enabling different forms of LTP to occur within the same neuron^24^, suggesting spatially heterogeneous calcium changes could mediate compartment-specific feedback. Consistent with this, regionally distinct calcium signals have been observed experimentally, including spatially restricted NMDA spike/plateau potentials confined to individual dendritic branches^25^.

Hence, we hypothesized that multiple, spatially segregated calcium signaling domains exist inside a neuron, enabling channel densities to be adjusted independently in different compartments (e.g. dendrite vs. soma). We tested our hypothesis in an abstract model and in a biophysically realistic multicompartment model. We found that dendritic and somatic calcium signals can remain spatially segregated, enabling the input-output relationship of separate compartments to be regulated independently. Our results also demonstrate (1) that homeostatic regulation based on somatic calcium leads to compensatory changes outside the perturbed region, (2) that compromising the spatial segregation of calcium signals diminishes compartment-specific regulation, and (3) that local perturbations exceeding the capacity of local compensation lead to neuron-wide homeostatic changes. These theoretical insights are crucial for appreciating the requirements and functional benefits of compartment-specific regulation, and for refining future experiments to delineate if/how such regulation is achieved and if it becomes pathologically altered.

## RESULTS

### Implications of local vs global feedback on the pattern of compensation in an abstract model

To explore the implications of controlling homeostatic regulation using local vs global feedback, we conducted exploratory simulations in a simple model. In this non-spiking model, input (i) is converted to output (o) based on a sigmoidal i-o relationship with adjustable gain. Transient variations in input produce transient variations in output but, if the input distribution were to experience a chronic shift, the output distribution can be restored by adjusting the gain (**Fig. 1A**); this equates in neurobiological terms to adjusting the synaptic weight or intrinsic excitability to restore the average firing rate to its target despite a sustained change in input. To incorporate two homeostatic mechanisms, the model was split into two compartments, where “synaptic scaling” controls the dendritic i-o relationship, or gain, and “excitability regulation” controls the somatic i-o relationship (**Fig. 1B**). Dendritic output constitutes the somatic input. If output deviates above or below its target, an error signal arises, triggering compensatory gain changes that minimize the error signal and, in the process, restores the output to its target. Though not applicable to our simple model, the error signal (calcium) is a proxy for the output (firing rate) in biophysical terms, opening up opportunities for misalignment; for example, direct perturbation of the error signal (not via the output signal) would cause inappropriate changes in the output^26^. Crosstalk between error signals could also distort compensation.

**Figure 1.**
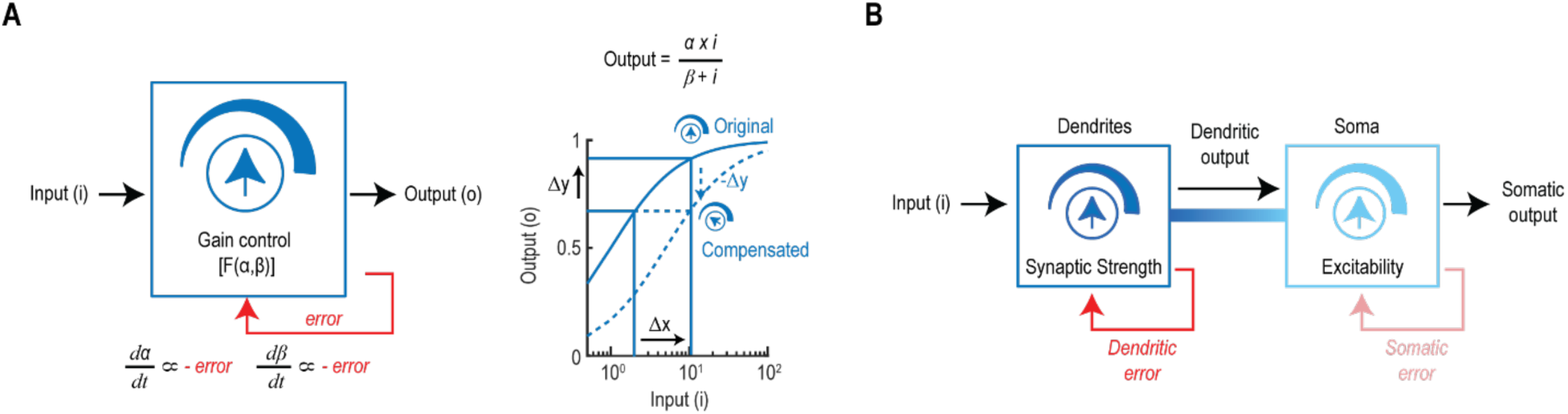
– An abstract non-spiking neuron model **A.** Left. Schematic of an abstract non-spiking model whose input-output relationship depends on the gain, which is a function of *α* and *β*, which are themselves adjusted based on an error signal (red). Right. A perturbation Δx in the input causes output to deviate Δy from its target value, producing an error signal. The error can be nullified by adjusting the gain (dashed curve). **B.** Variation of the model in A comprising a dendritic and somatic compartment. The dendritic compartment receives input that is transformed into dendritic output based on synaptic strength (i.e. dendritic gain). The somatic compartment receives dendritic output as input and transforms it into somatic output based on its intrinsic excitability (i.e. somatic gain). An important consideration is whether the gain of each compartment is independently regulated using separate error signals (as illustrated here), which would mediate local homeostatic changes; a shared error signal is predicted to produce non-local homeostatic changes.

First, we simulated the pattern of compensatory changes induced by a local perturbation when feedback is also local (**Fig. 2**). When a sustained increase in input was applied to the dendritic compartment (**Fig. 2A top**), the dendritic output and error signal were both initially increased, but both returned to baseline because of changes in the dendritic gain, manifest as sustained changes in parameters *α* and *β*. The soma exhibited a transient increase in its output and local error signal, but the change in its gain was transient, unlike the sustained change in dendritic gain. Ultimately, the somatic output was restored to baseline by the restoration of somatic input via changes in dendritic gain, which rendered downstream changes in the somatic gain unnecessary. In other words, a perturbation to the dendrites was corrected entirely by homeostatic changes in the dendrites. The same pattern of sustained changes in dendritic gain and transient changes in somatic gain was observed during a sustained decrease in dendritic input (**Fig. 2A bottom**).

**Figure 2.**
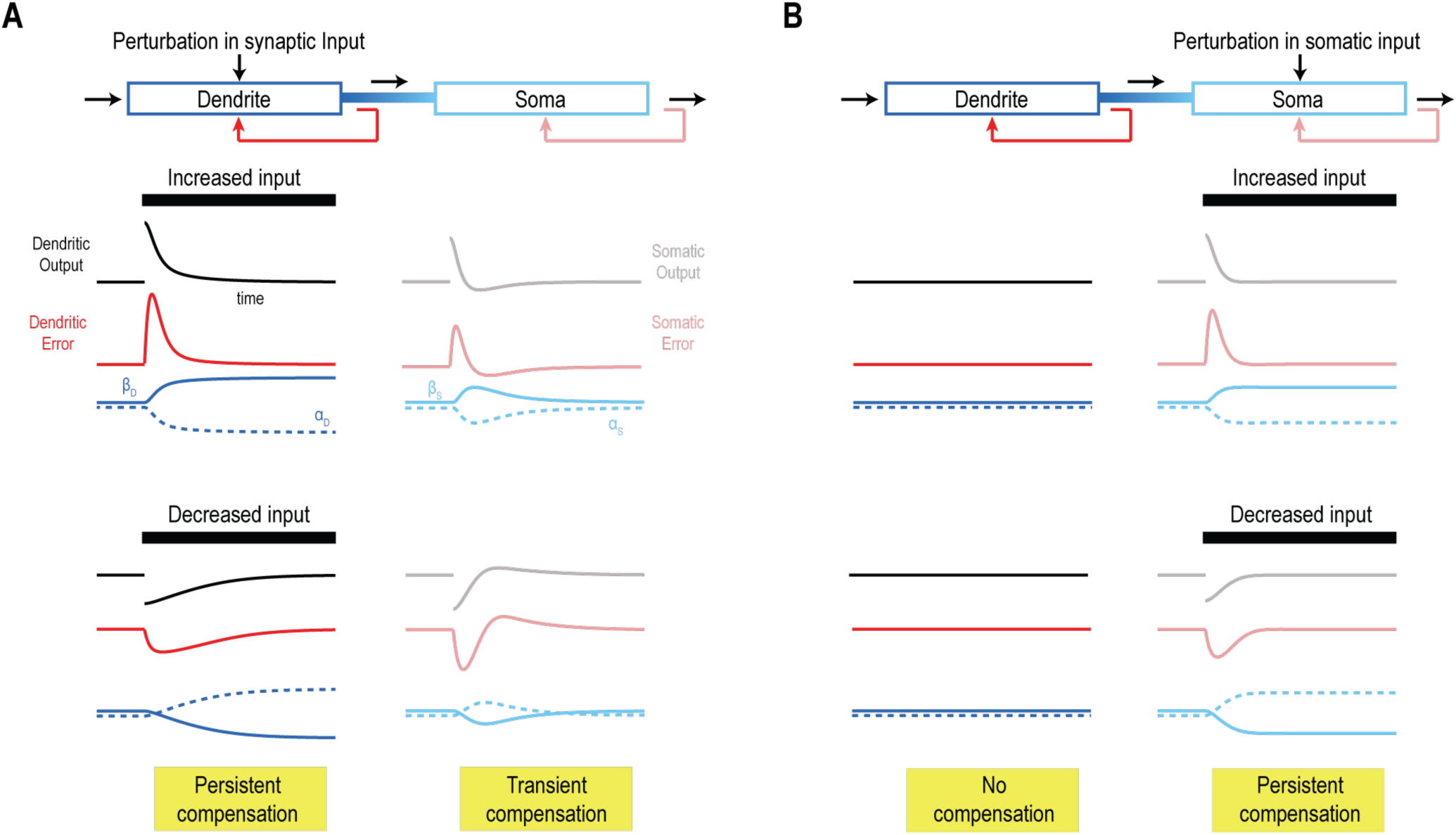
– Abstract model with local (compartment-specific) feedback exhibits sustained homeostatic changes only in the perturbed compartment. Output (black), error (red), and gain control parameters (blue; α, dashed; β, solid) are shown for the dendrites (left, dark) and soma (right, pale). **A.** Dendritic perturbations. For an increase in input (top), both compartments exhibit a transient change in output that is paralleled by the local error signal. Both compartments initially exhibit changes in parameters *α* and *β*, but those changes are only sustained in the dendrites, demonstrating that when dendritic output is corrected (by adjusting dendritic gain), compensation is no longer required in the soma. The same pattern is observed during a decrease in input to the dendrites (bottom). **B.** Somatic perturbations. Because of the unidirectional flow of information, only the somatic output is altered when somatic input is perturbed. But like in A, the compartment experiencing the perturbation (in this case the soma) exhibits a sustained change in parameters *α* and *β* so that its output is corrected.

Next, we simulated a sustained increase in somatic input (**Fig. 2B top**). Like in panel A, the compartment experiencing the perturbation, in this case the soma, exhibited only a transient increase in its output and error signal; both were returned to baseline by sustained changes in somatic gain. Unlike in panel A, the other compartment, namely the dendrites, did not experience any changes because the flow of information is unidirectional from dendrites to soma (and because there is no crosstalk between dendritic and somatic feedback). The same compensation pattern was observed during a sustained decrease in somatic input (**Fig. 2B bottom**).

Results in Figure 2 establish that effects of a perturbation to one compartment can be offset by compensatory changes in that compartment when feedback is local. Though rather obvious, this nevertheless serves as an important comparison for what happens when feedback is not compartment-specific, which is what we simulated next, namely global feedback.

In the same model but now using a single error signal that represents combined dendritic and somatic errors, an increase in dendritic input (**Fig. 3A top**) was found to cause sustained changes in both the dendritic and somatic gain. The somatic output and the global error signal both returned to baseline, but the dendritic output remained elevated; this occurs because compensatory gain changes are distributed between two compartments, which means the dendritic output, although partially compensated, remains elevated to accommodate the change in somatic gain. The same pattern was observed in response to a sustained decrease in dendritic input (**Fig. 3A bottom**). A similar pattern was also observed during a sustained increase in somatic input (**Fig. 3B top**). In this case, the somatic output and global feedback signal both returned to baseline thanks to combined compensatory changes in dendritic and somatic gain, but this introduced a sustained decrease in dendritic output. The same pattern was observed in response to a sustained decrease in somatic input (**Fig. 3B bottom**).

**Figure 3.**
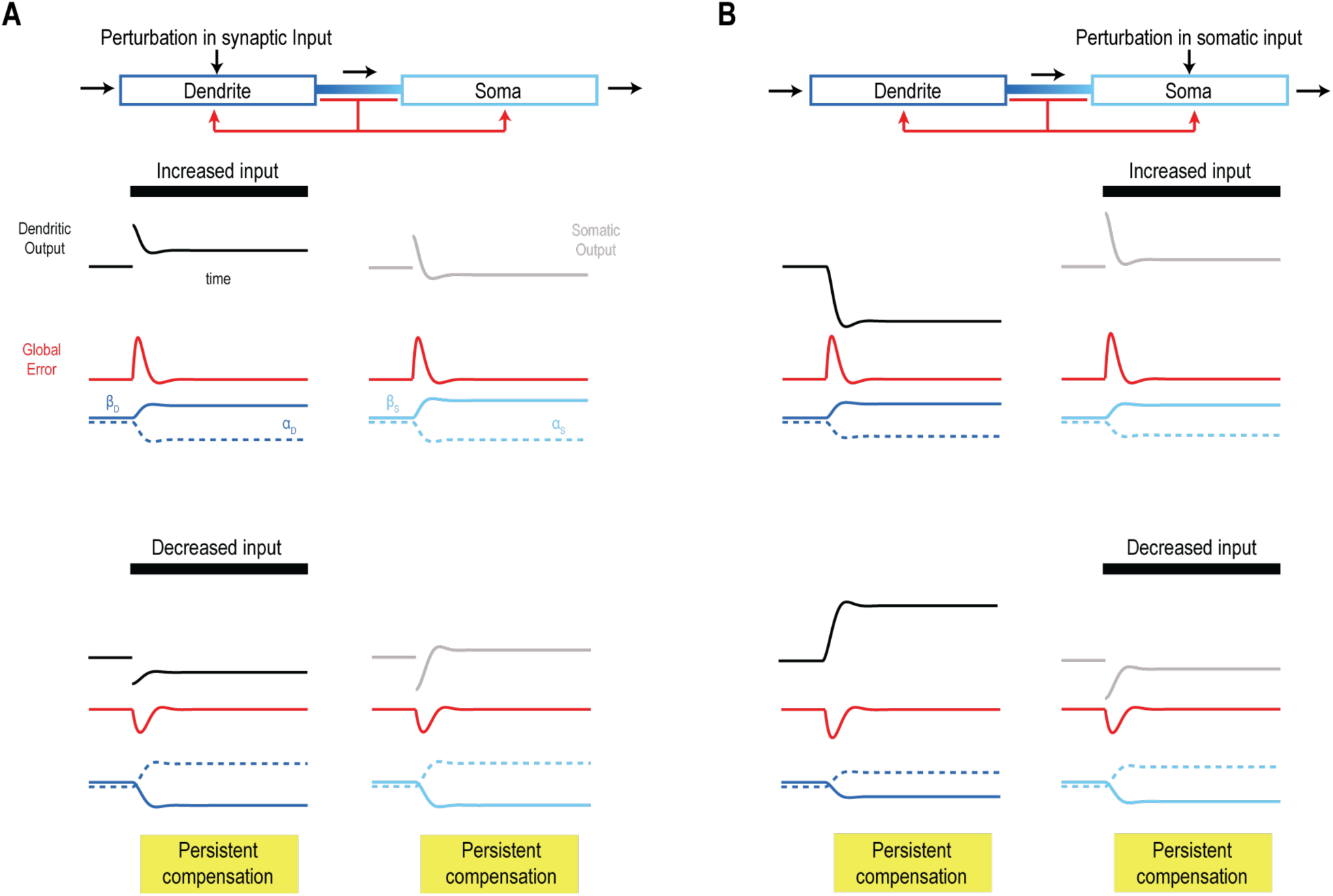
– Abstract model with global feedback exhibits sustained homeostatic changes in both compartments. Output (black), error (red), and gain control parameters (blue; α, dashed; β, solid) are shown for the dendrites (left, dark) and soma (right, pale). **A.** Dendritic perturbations. During a sustained increase (top) or decrease (bottom) in dendritic input, both compartments exhibit a sustained change in their gain, unlike in Figure 2. Related to this, dendritic output is only partially compensated because compensation also occurs in the soma (because of the shared error signal), which means compensation occurs in both compartments regardless of which one is perturbed. **B.** Somatic perturbations. A pattern of distributed compensation similar to panel A is observed. Notably, the global error signal induces compensation upstream of the perturbation, which leads to a change in dendritic output, which partially offsets the perturbation to somatic input. In short, compensatory changes are not limited to the compartment experiencing the perturbation.

Results in Figure 3 establish that global feedback leads to distributed compensatory changes in gain. In other words, a perturbation applied to one compartment does not trigger compensatory changes exclusive to that compartment (even when the two compartments are unidirectionally connected) because the error signal spans both compartments. Instead, distributed compensation may only partially restore a perturbed local i-o relationship (as in Fig. 3A) and may even change a local i-o relationship that was not *directly* impacted by the perturbation (as in Fig. 3B). In other words, non-local feedback leads to non-local compensation. In this case, the global i-o relationship (the relationship between dendritic input and somatic output) is restored regardless of where the perturbation occurs, but local i-o relationships (dendritic i-o and somatic i-o) are not returned to normal, which could have important functional consequences.

We predicted that applying a perturbation to one dendrite would alter the i-o relationship of a second dendrite if feedback is global and, consequently, compensation is distributed. Specifically, we predicted that the relationship between input to the perturbed dendrite and somatic output driven by that input would benefit from the compensation, but that the relationship between input to the other dendritic and somatic output would suffer from “off-target” compensation, which is relevant for any scenario where multiple dendrites converge on a common soma. To test this, we added a second dendritic compartment to the abstract model, where both dendrites are unidirectionally connected to the soma. As predicted for local feedback, when a sustained increased input was applied to dendrite 1 (**Fig. 4A**), its output and error signal were both initially increased but both returned to baseline because of compensatory changes in gain; the soma exhibited transient changes (like in Fig. 2A) but dendrite 2 was unaffected. Perturbing the somatic input led to changes exclusively in the soma (**Fig. 4B**; like in Fig. 2B). With global feedback, on the other hand, increased input to dendrite 1 led to a compensatory reduction in gain across both dendrites and the soma (**Fig. 4C**), which is reminiscent of changes in Figure 3A, where increased output from the perturbed compartment is only partially reduced because of changes occurring in other compartments. Notably, the output of dendrite 2 was reduced below baseline. Like in Figure 3B, an increase in somatic input led to reduced gain in both dendrites (**Fig. 4D**). Therefore, as predicted, global feedback leads to compensation in unperturbed compartments, inadvertently altering their input-output relationship while blunting compensation in the perturbed compartment.

**Figure 4.**
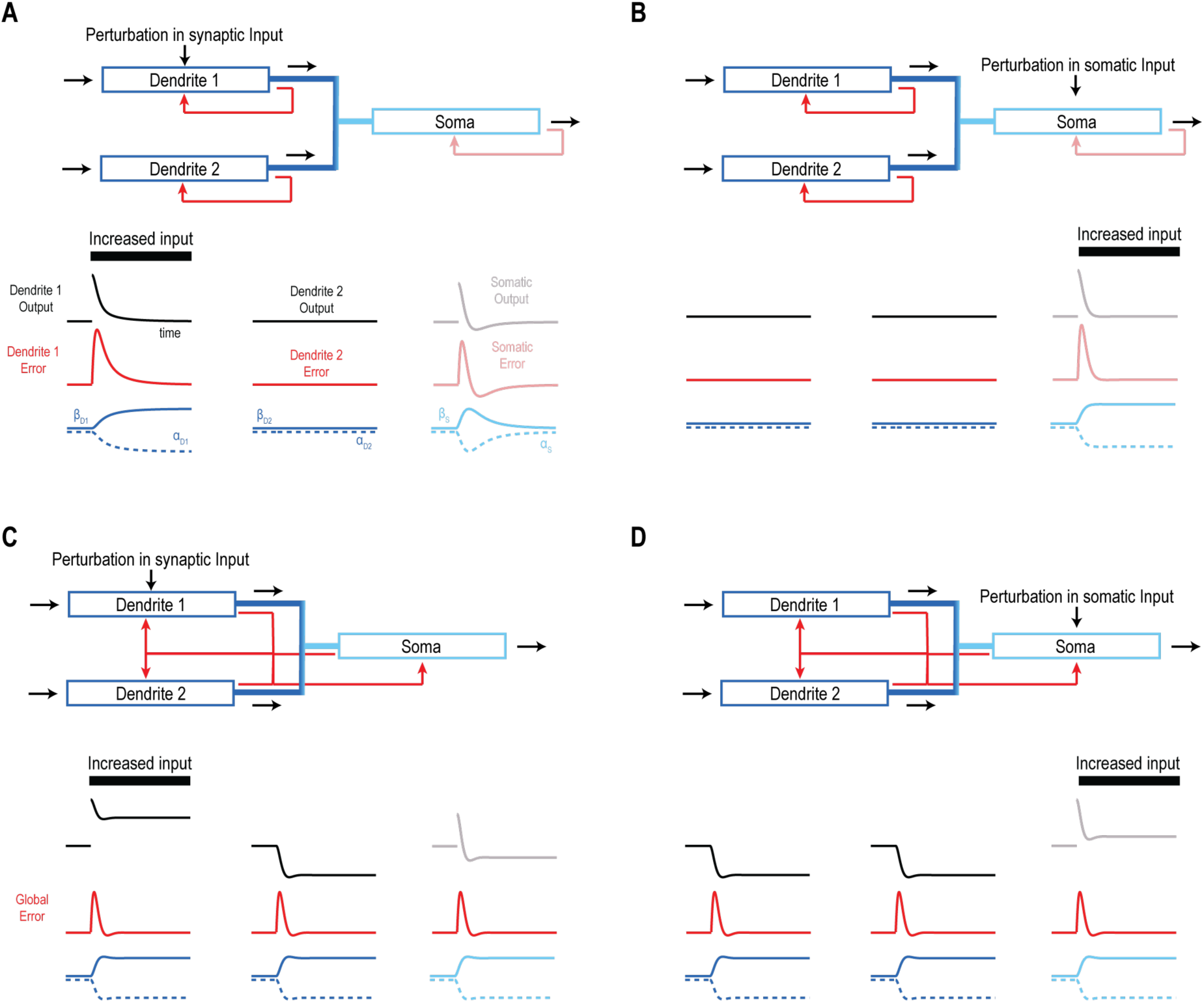
– Abstract model with three compartments and global feedback exhibits homeostatic changes in unperturbed compartments. Output (black), error (red), and gain control parameters (blue; α, dashed; β, solid) are shown for the dendrites (left and center, dark) and soma (right, pale). **A.** Increased input to dendrite 1 with local feedback. Dendrite 1 and the soma exhibit a transient change that is paralleled by the local error signal; both compartments initially exhibit changes in parameters *α* and *β*, but those changes are only sustained in dendrite 1. Dendrite 2 does not exhibit any effects since information flows unidirectionally from dendrites to soma and because feedback is local. **B.** Increased input to soma with local feedback. Because of the unidirectional flow of information, only the somatic output is altered. Like in panel A, only the compartment experiencing the perturbation (in this case the soma) exhibits a sustained change in parameters *α* and *β* so that its output is corrected. **C.** Increased input to dendrite 1 with global feedback. All compartments exhibit a sustained change in their gain, which is unlike in panels A and B. Related to this, dendrite 1 output is only partially compensated because compensatory changes in parameters *α* and *β* also occur in dendrite 2 and the soma (because of the shared error signal), which means compensation occurs in all compartments regardless of which one is perturbed. **D.** Increased input to soma with global feedback. A pattern of distributed compensation similar to panel C is observed. Notably, the global error signal induces compensation upstream of the perturbation, which leads to a dramatic change in both dendritic outputs which then partially offset the perturbation to somatic input. In short, compensatory changes are not limited to the compartment experiencing the perturbation.

#### Realistic neurons can maintain spatially segregated calcium domains despite diffusion

Results from the abstract model provide proof-of-principle evidence that global feedback leads to a pattern of distributed compensation that is distinct from the more targeted compensation enabled by local feedback. However, this raises the question of whether the intracellular calcium changes thought to encode error signals can achieve the spatial segregation required to mediate feedback independently in each compartment. There are several considerations. First, calcium influx occurs through both synaptic receptors and voltage-gated calcium channels (VGCCs); the former are activated by synaptic input whereas the latter are activated primarily by spiking (insofar as high-voltage-activated calcium channels are activated mostly, if not exclusively, during the strong depolarization associated with a spike). And while most excitatory synapses exist in the dendrites (and thus affect dendritic calcium more than somatic calcium), most spikes originate in/near the soma but can backpropagate into the dendrites, thus causing calcium influx via VGCCs in both the soma and dendrites. Furthermore, calcium diffuses intracellularly between compartments. Thus, it is not obvious that dendritic and somatic calcium provide independent, spatially restricted error signals suitable for controlling synaptic scaling and excitability regulation, respectively. Moreover, it arguably make sense to treat excitability regulation in the soma and dendrites as separate processes since the latter contributes (along with synaptic scaling) to dendritic gain. Thus, to explore the biological plausibility of independent calcium-based feedback signals, we built a biophysically realistic multicompartment spiking model in which to run tests akin to those conducted in our abstract model.

We used the morphology of a reconstructed pyramidal neuron^27,28^ with synapses throughout the dendrites and voltage-gated ion channels distributed according to reported patterns^29,30^ (**Fig. 5A**); densities were adjusted to reproduce numerous response properties (see Methods). With parameters thus set, the model was exposed to two perturbations. In the first, increasing synaptic input to three oblique dendrites elicited dendritic spikes that were attenuated to small EPSPs upon reaching the soma (**Fig. 5B left**), consistent with past experimental data^31^. In the second perturbation, increasing current injection at the soma elicited action potentials that propagated back into distal dendrites but with decreasing amplitude (**Fig. 5B right**), consistent with experimental data^30,32^. The first perturbation (to three oblique dendrites) caused a calcium increase locally that was about 10x larger than in the apical dendrite or soma (**Fig. 5C**). The second perturbation (to the soma) caused an increase in somatic calcium that spread into the apical dendrite primarily because of bAPs, but the somatic increase was still ∼3x larger than in oblique dendrites targeted in the first perturbation (**Fig. 5D**). These data show that calcium changes are largest near the perturbation, indicating some degree of segregation.

**Figure 5.**
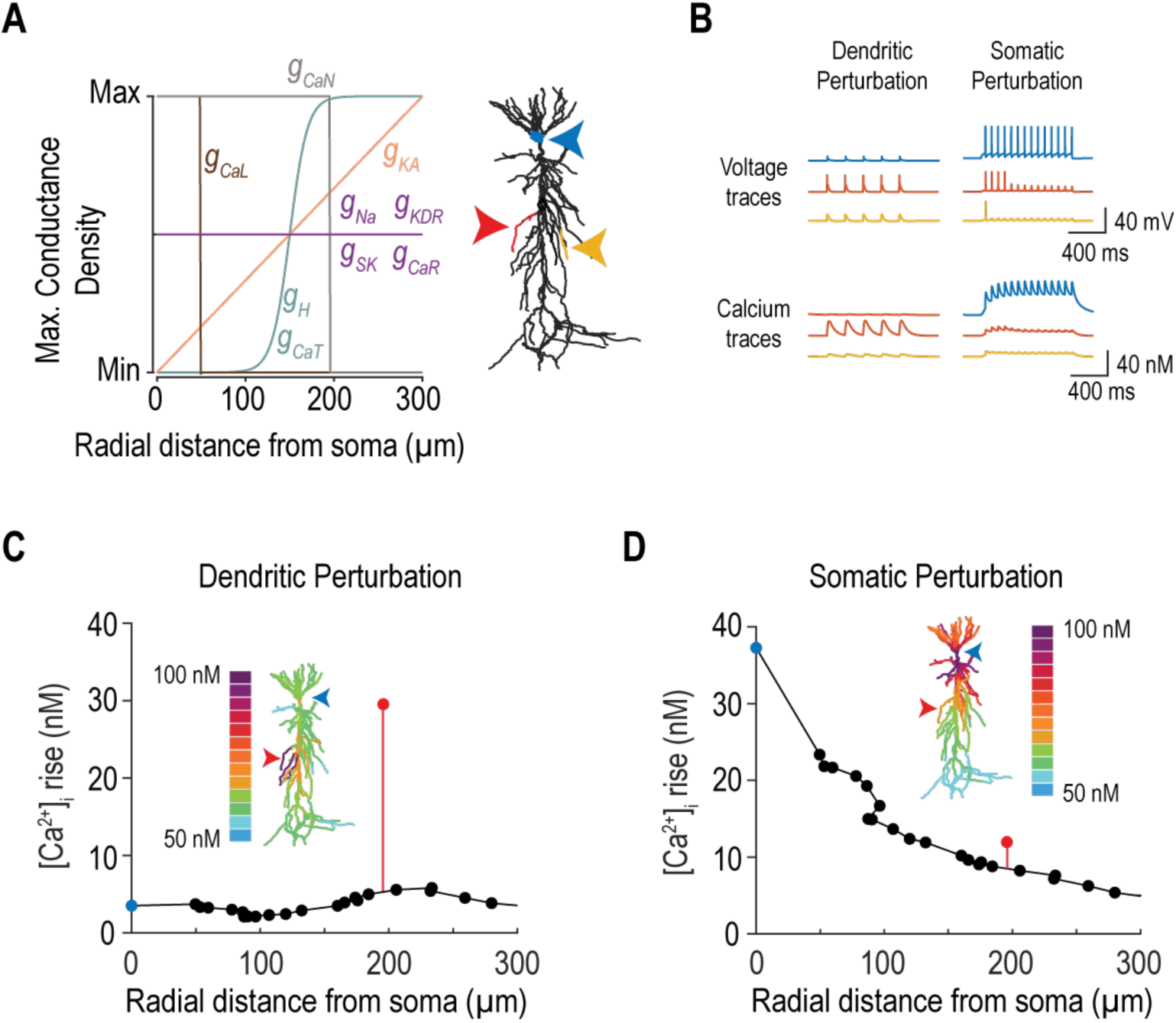
– Realistic spiking model exhibits segregated calcium changes that could support local feedback. A. Distribution of ion channels as a function of radial distance from the soma. Inset shows morphology of the multicompartment model based on a reconstructed pyramidal neuron; colored arrows show recording sites for traces in panel B. B. Left. Sample traces for voltage and intracellular calcium concentration [Ca²⁺]_i_ illustrate that a dendritic perturbation (increased synaptic input) leads to local dendritic spikes and significant local calcium influx (red traces), while only small amplitude synaptic potentials reach the soma (blue trace) and non-stimulated dendritic branches (yellow trace), both of which therefore experience little change in calcium. Right. Sample traces show that action potentials initiated at the soma (blue trace) due to a somatic perturbation (depolarizing current injection) propagate into the dendrites but fail to reach distal dendrites (red and yellow traces), limiting the calcium influx experienced in distal dendritic regions. C. Calcium changes due to a perturbation in a set of oblique dendrites (same as panel B, left), plotted along the apical dendrite (black dots) and at the soma (blue) and perturbed oblique dendrite (red) confirm that the elevation in [Ca²⁺]_i_ is limited to the perturbed dendrite. Inset shows [Ca²⁺]_i_ at steady state. D. Calcium changes due to a somatic perturbation (same as panel B, right) spread into the dendrites but are nonetheless attenuated over space.

#### Differential calcium changes enable independent regulation in the dendrites and soma

Next, we checked whether the relatively localized calcium changes reported in Figure 5 could support independent homeostatic regulation in the dendrites and soma. The strength of AMPAR and NMDAR synapses and the densities of fast sodium and M-type potassium channels were adjusted using a local calcium-based error signal (see Methods). Four different perturbations were tested. When a sustained increase in input was applied to the oblique dendrites (**Fig. 6A**), the model neuron exhibited an initial increase in firing rate (i), which included more dendritic spikes, but firing rate eventually returned to baseline. Changes in calcium (ii) were also transient, with the largest change occurring in the perturbed dendrite. The compensatory change in *g̅*_Na_(iii) was sustained in the perturbed dendrite but transient in the soma. The compensatory change in *g̅*_KM_(iv), which exists uniquely in the soma, was also transient. The compensatory change in *g̅*_AMPA_ (v) was sustained in the perturbed dendrite but transient in a nearby dendrite. The same pattern was observed in response to a sustained reduction in dendritic input (**Fig. S1A**). This pattern is very similar to that seen in the abstract model with local feedback (see Fig. 2); specifically, once the effect of the perturbation was offset by local compensatory changes, compensatory effects that had started to develop elsewhere waned. A similar pattern of local compensation was observed in the soma when its input was increased (**Fig. 6B**) or decreased (**Fig. S1B**); specifically, compensatory changes in the soma were sustained whereas compensatory changes in the dendrites were transient. Dendrites and soma are bidirectionally coupled in the realistic model, unlike in the abstract model.

**Figure 6.**
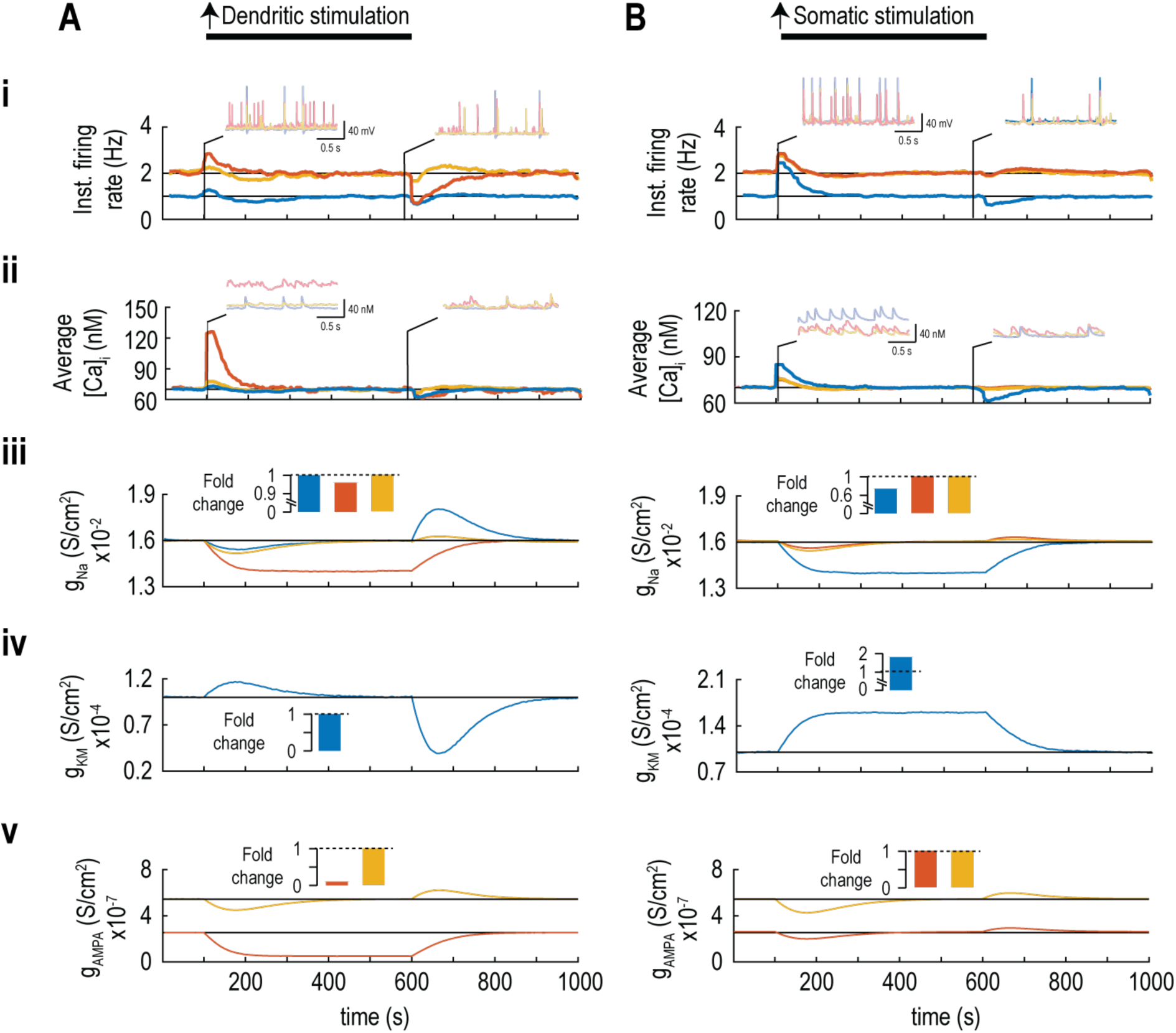
– Realistic spiking model can exhibit localized homeostatic changes in response to local perturbations. Traces are color coded based on recording site (see Fig. 5A). All firing rate and intracellular calcium traces are smoothened using a 20s rolling average filter. **A.** Increased dendritic stimulation. Firing rate (i) and intracellular calcium (ii) were altered by the perturbation but returned to baseline following compensatory changes (iii-v). Top insets show sample voltage and calcium traces at two time points indicated by black lines. Compensatory changes in *g̅*_Na_(iii), *g̅*_KM_(iv), and *g̅*_AMPA_ (v) were transient in all compartments except in the perturbed dendrite where changes were sustained. Bar graph insets summarize the fold change in conductance densities at steady state (i.e. just prior to termination of the perturbation). **B.** Increased somatic stimulation. Data are plotted like in A and reveal equivalent results except that sustained compensation occurs uniquely at the soma (blue), which is subject to perturbation.

Notably, the calcium increase at the soma caused by a somatic perturbation (**Fig. 6Bii**) was larger than in the dendrites; by comparison, the calcium increase in oblique dendrites experiencing the dendritic perturbation was much larger than in the soma (**Fig. 6Aii**), consistent with Figure 5. An important consideration is whether remote calcium changes occur secondary to the local calcium change driven directly by a perturbation. If compensation is controlled by the local calcium change, then local compensation should nullify the primary calcium change and, in turn, secondary calcium changes should wane; for example, if compensatory changes in somatic excitability nullify the effects of a somatic perturbation on firing rate, the disturbance in dendritic calcium due to bAPs will also return to normal without any sustained compensatory change in the dendrites. However, if compensation is controlled by a remote increase in calcium – for simplicity, imagine dendritic compensation being controlled by somatic calcium changes – then somatic compensation is liable to nullify the change in somatic calcium faster or more effectively than dendritic compensation even if the perturbation occurs in the dendrites. To test this, we repeated simulations with an increase in dendritic input (like in Fig. 6A) but we made all compensation dependent on somatic calcium. We predicted that compensation would be distributed between the soma and dendrites, like in Fig. 3A and unlike Figs. 2A and 6A. **Figure 7A** confirms that if dendritic compensation depends on changes in somatic calcium, dendritic perturbations lead to sustained compensation across the neuron, including in the soma. Moreover, rectifying a calcium change in the soma might not completely rectify dendritic calcium changes, allowing changes in the input-output relationship between dendrites and soma to go uncorrected.

**Figure 7.**
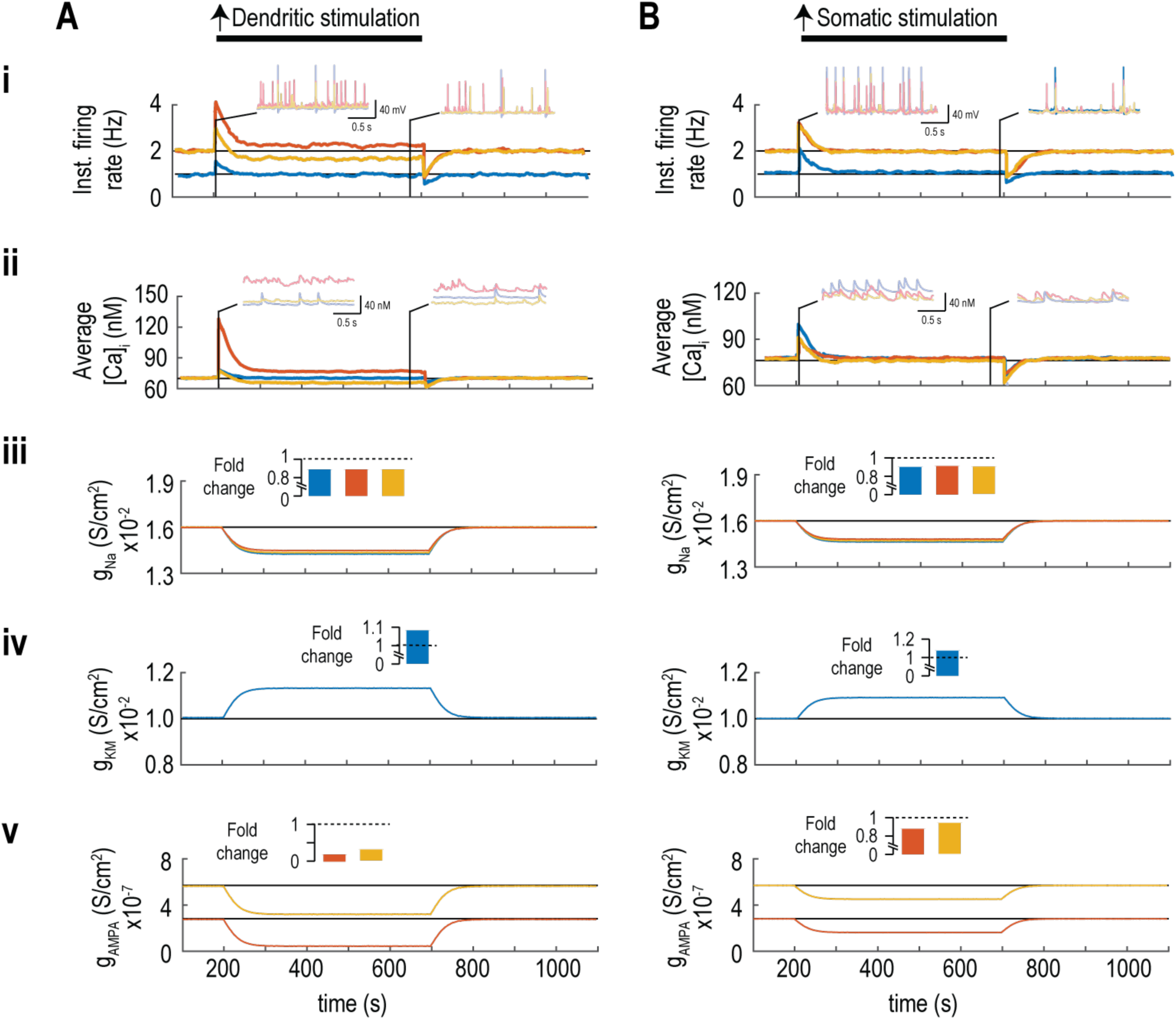
– Realistic spiking model with compensation controlled by somatic calcium exhibits incomplete off-target compensation. Traces are color coded based on recording site (see Fig. 5A). All firing rate and intracellular calcium traces are smoothened using a 20s rolling average filter. **A.** Increased dendritic stimulation. Firing rate (i) and intracellular calcium (ii) were altered by the perturbation but compensation returned them to baseline only in the soma. Overcompensation of firing rate and calcium is seen for the unperturbed dendrite (yellow) and under-compensation is observed for the perturbed dendrite (red). Compensatory changes in *g̅*_Na_ (iii), *g̅*_KM_ (iv), and *g̅*_AMPA_ (v) were sustained in all compartments. Sample traces are shown for two time points indicated by black lines. Bar graph insets summarize fold changes in conductance densities at steady state (i.e. just prior to termination of the perturbation). **B.** Increased somatic stimulation. Data (plotted like in A) reveal that the firing rate and calcium in each compartment was brought back to normal with sustained compensatory changes in *g̅*_Na_, *g̅*_KM_, and *g̅*_AMPA_ occurring in all compartments.

We also repeated simulations with an increase in somatic input (like in Fig. 6B) but again with all compensation dependent on somatic calcium. We once again predicted that compensation would be distributed across the soma and dendrites, like in Fig. 3B and unlike Figs. 2B and 6B. **Figure 7B** confirms that if neuron-wide compensation depends on somatic calcium changes, somatic perturbations lead to sustained compensation across the neuron, including in dendrites.

#### Increased calcium crosstalk reduces the independence of homeostatic regulation

Next, we investigated how reducing the spatial segregation of calcium signals impacts local homeostatic changes. This question is relevant in certain neurodevelopmental disorders where reduction of A-type potassium channels in dendrites may reduce the attenuation of bAPs^33^. An increase in bAP amplitude would increase calcium influx in distal dendrites due to enhanced activation of high-voltage-activated calcium channels, thereby diminishing the spatial segregation between somatic and distal dendritic calcium signals. To test this, the distribution of A-type potassium channels, whose density in the dendrites normally increases with distance from the soma (see Fig. 5A), was changed to a uniform distribution. We re-tuned other parameters to restore the average firing rate and calcium levels to values seen in the original model (see Table 3) but the change in A-type channel distribution nevertheless increased the spread of active signals toward or away from the soma (depending on the stimulation site), which decreased calcium segregation, as demonstrated below using perturbations implemented like in Figures 5 and 6.

In the revised model, diminished attenuation of dendritic spikes evoked by stimulating a set of oblique dendrites (**Fig. 8Ai top**) resulted in larger calcium changes in unperturbed compartments (**Fig. 8Ai bottom**). This translates to less spatially segregated calcium changes (**Fig. 8Aii**) which, in turn, led to persistent compensatory changes in *g̅*_Na_, *g̅*_AMPA_, and *g̅*_KM_ that were non-local (i.e. in the soma and/or unperturbed dendrites), unlike in the original model (**Fig. 8Aiii**; see also **Fig. S2A**). During somatic stimulation, a similar reduction in the attenuation of bAPs was observed, which again led to less spatially segregated calcium changes and non-local compensatory changes (**Figs. 8B** and **S2B**). In short, a change in the distribution of A-type potassium channels, despite its most obvious effects on output being nullified by adjustments in other channels, led to increased calcium cross-talk leading to non-local compensation.

**Figure 8.**
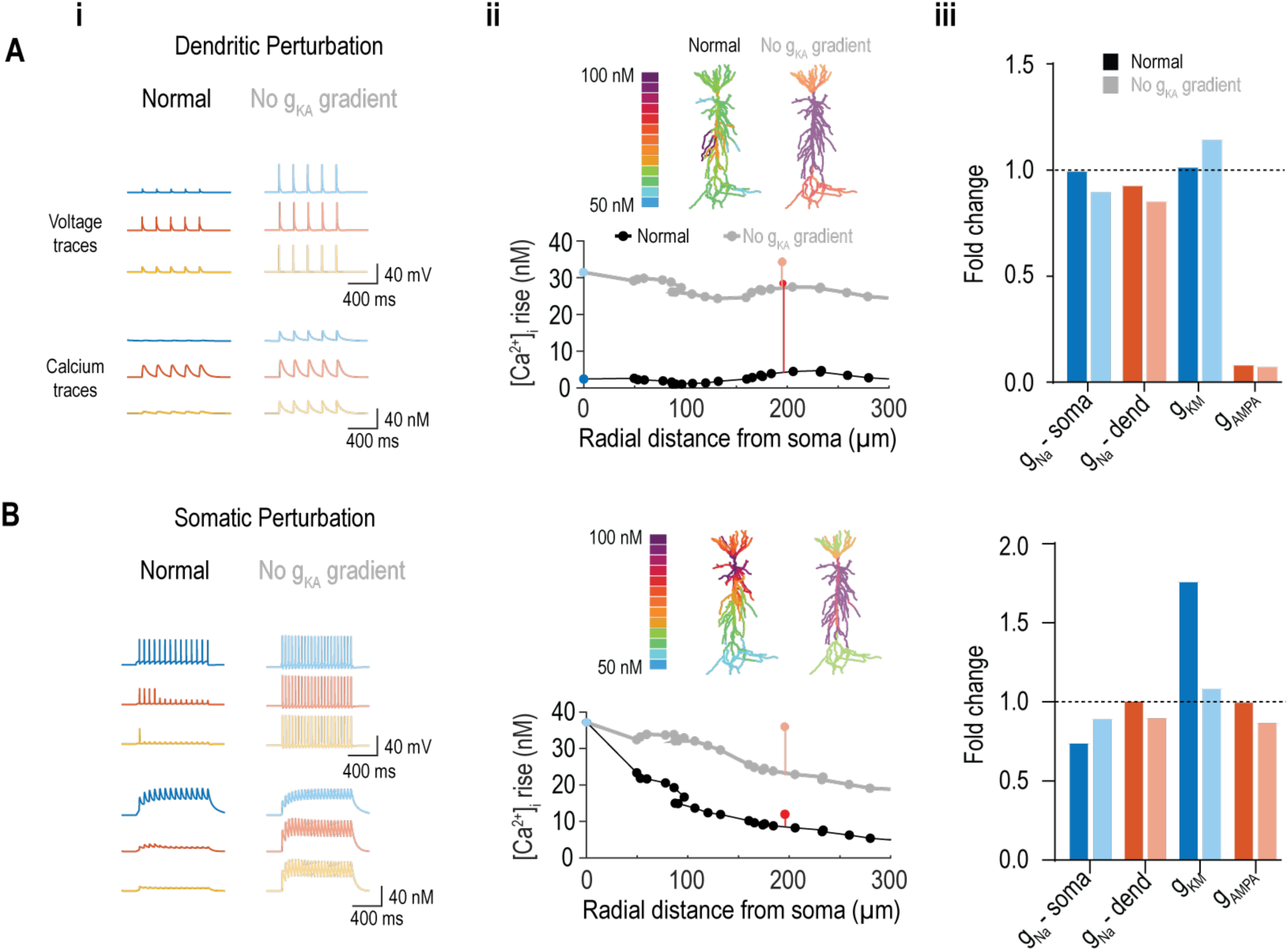
– Modified model with uniformly distributed A-type potassium channels exhibited less segregated calcium signaling, resulting in non-local homeostatic changes. See Fig. S2 for a full set of sample responses in the modified model. **A.** Increased dendritic stimulation. (i) In contrast to the original model (left), dendritic spikes more efficiently propagated toward the soma to trigger somatic spikes, and also spread to other dendrites, resulting in a broader increase in calcium in the modified model (right). (ii) The peak rise in [Ca²⁺]_i_ documents the broader increase in calcium that results, which is also evident in the steady state [Ca²⁺]_i_ (insets). (iii) Compared to the compensation pattern in the original model (dark bars), the pattern of homeostatic changes in the modified model (pale bars) involves a greater-than-expected reduction in somatic *g̅*_KM_and a smaller-than-expected reduction in somatic *g̅*_Na_, coupled with diminished compensation in the dendrites, highlighting the shift to non-local compensatory changes. **B.** Increased somatic stimulation. (i) Compared to the original model (left), somatic spikes propagated less attenuated into the distal dendrites in the modified model (right), causing increased calcium influx distally. (ii) Bottom graph and insets document the broader changes in [Ca²⁺]_i_.. (iii) The altered compensation pattern demonstrates non-local decreases in dendritic *g̅*_Na_and *g̅*_AMPA_coupled with diminished compensation in the soma, particularly for *g̅*_KM_.

#### Excessively strong perturbations lead to global compensation

Pharmacological agents like tetrodotoxin (TTX) are frequently used in experiments to trigger homeostatic changes^3,11,34,35^. Such an intervention constitutes an incompensable perturbation insofar as neurons cannot restore spiking in the presence of TTX. We hypothesized that local incompensable perturbations will saturate local compensatory mechanisms and cause compensation to "spillover" to other parts of the neuron, for instance, causing synaptic upscaling in an effort to restore spiking if increasing intrinsic excitability cannot restore spiking. To test this, we modeled blockade of spiking within 100 µm of the soma, effectively abolishing somatic spikes (**Fig. 9i**, blue) and significantly lowering somatic calcium levels (**Fig. 9ii**, blue). This perturbation elicited saturated compensatory responses, notably in somatic *g̅*_KM_ (**Fig. 9iv**), causing excessive compensation in dendritic firing rates (**Fig. 9i**, red and yellow) and calcium (**Fig. 9ii**, red and yellow). Conductance changes included pronounced and sustained increases in somatic *g̅*_Na_ (**Fig. 9iii**, blue), dendritic *g̅*_Na_ (**Fig. 9iii**, red and yellow), and dendritic *g̅*_AMPA_ (**Fig. 9v**). These findings suggest that intense perturbations exceeding local compensatory capacity trigger homeostatic adjustments across the entire neuron, even when perturbations (and error signals) are local. This finding suggests that local compensation may not have been observed experimentally because testing has typically involved incompensable perturbations.

**Figure 9.**
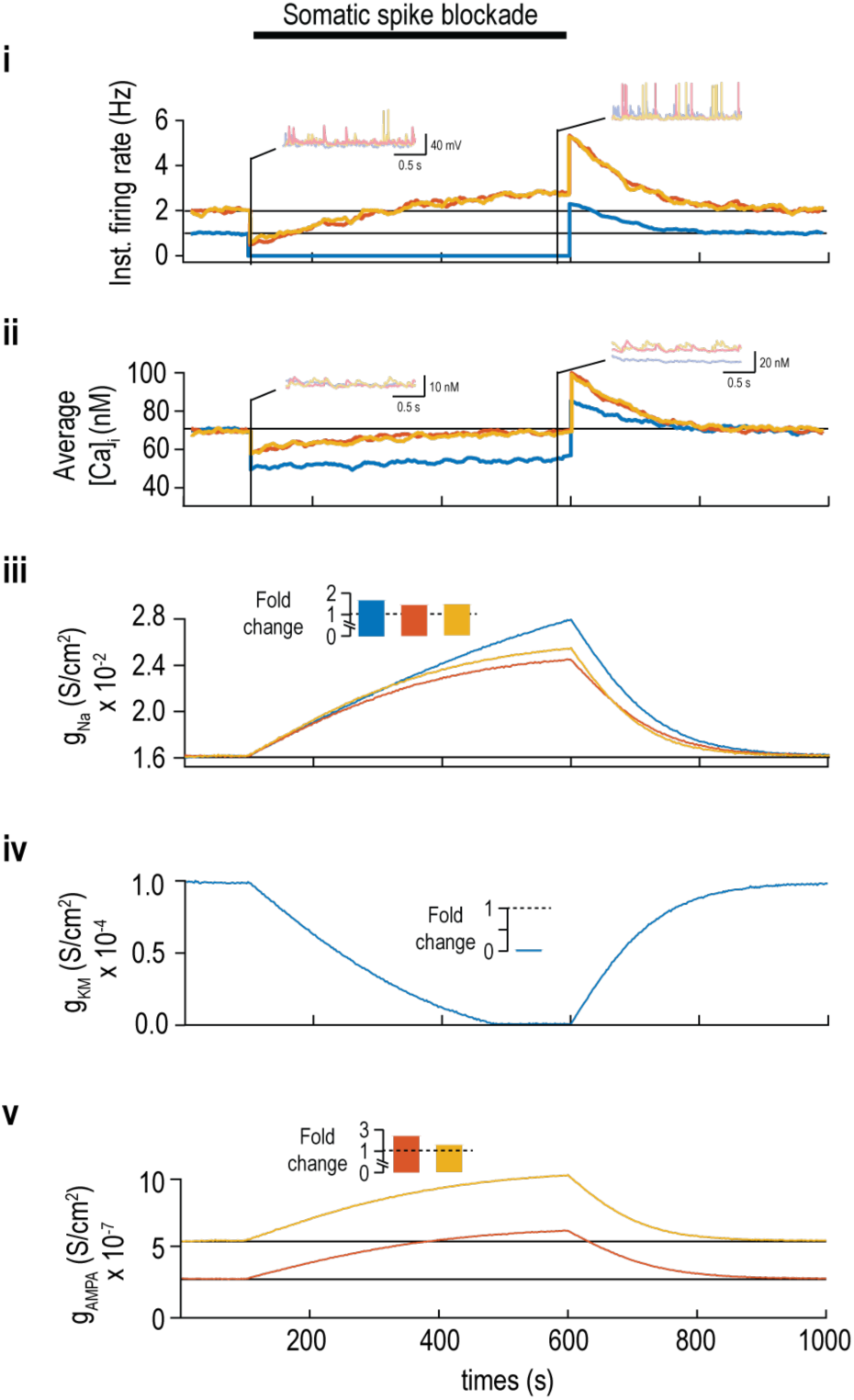
– Realistic spiking model experiencing an incompensable perturbation exhibits non-local compensation. During blockade of spikes in the soma, local compensatory changes (directed by a local error signal) could not overcome the blockade to restore somatic firing rate (i) and could not, therefore, restore somatic calcium (ii) despite large changes in *g̅*_Na_(iii) and *g̅*_KM_ (iv). This led to compensatory changes beyond the soma, including changes in *g̅*_AMPA_ (v). Compensation in the dendrites led to overcorrection of dendritic output. All firing rate and intracellular calcium traces are smoothened using a 20s rolling average filter.

## Discussion

Starting with a simple two-compartment model, we demonstrated that if activity-dependent feedback operates independently in each compartment (using a local error signal), then compensatory changes persist exclusively in the perturbed compartment (Fig. 2). Compensatory changes initially develop in other compartments but reverse once the input-output relationship of the perturbed compartment is normalized. In contrast, if compensation relies on shared feedback (using a non-local error signal), perturbing one compartment leads to other compartments experiencing sustained compensatory changes (Figs. 3 and 4); in this scenario, the global input-output relationship is normalized but the input-output relationships of individual compartments become distorted. We then validated these observations in a morphologically and biophysically realistic neuron model. Despite calcium being able to diffuse within the cell, and despite calcium influx caused by spikes propagating bidirectionally through the dendrites, our realistic model revealed that calcium changes triggered by local perturbations remain relatively segregated (Fig. 5), thus mitigating cross-talk between error signals and enabling local homeostatic changes (Fig. 6). Our results show that homeostatic changes are non-local if feedback is based on somatic calcium (Fig. 7), if the segregation of calcium-based error signals is compromised (Fig. 8), or if a perturbation is too large to be offset by local compensation (Fig. 9).

In our two-compartment model (Fig 1B), regulation of dendritic and somatic gain were equated with synaptic scaling and excitability regulation, respectively. This was prompted by the notion that these two forms of homeostatic regulation might be independently regulated. But in our multicompartment model (Fig. 5A), dendrites were endowed with various voltage-gated channels that influence the spread of excitatory postsynaptic potentials (EPSPs) toward the soma; in other words, dendritic excitability – in addition to synaptic strength – shapes dendritic output. Accordingly, we modeled synaptic scaling and regulation of dendritic excitability as being controlled by the same local calcium signal, separate from excitability in the soma, which was controlled by a different local calcium signal. In that respect, feedback was independent between compartments, but synaptic scaling and dendritic excitability regulation were linked. They could be regulated independently if different error signals were involved, including error signals other than activity-dependent calcium changes, but we did not consider this in our simulations. Prior modeling likewise indicates that dendritic calcium sources can be organized into spatially segregated domains; discrete recruitment of dendritic CaV1.3 clusters in motoneurons, for example, implies multiple, separately addressable dendritic calcium compartments^36^.

Many studies have reported homeostatic regulation of synaptic strength and intrinsic excitability occurring concurrently. For instance, application of TTX to cultured cortical pyramidal neurons triggers a compensatory increase of both synaptic strength^37^ and intrinsic excitability^3^. Ibata et al.^11^ showed that blockade of postsynaptic spiking (by TTX) triggered synaptic upscaling whereas local blockade of synaptic input did not. Contrary to this, Fong et al.^38^ showed that synaptic upscaling was triggered by blocking synaptic input, even when spiking was restored through closed-loop optogenetic stimulation. Hyperpolarizing neurons by overexpressing potassium channels (rather than altogether blocking spikes with TTX) was shown to have different effects depending on the developmental status of tested neurons^39^. Interfering with AMPA receptor trafficking caused a compensatory increase in excitability^40^. The nature of the perturbation is evidently important, but most data suggest interdependent regulation of synaptic strength and excitability. Our results highlight the importance of perturbation intensity: non-local regulation occurs following strong perturbations not necessarily because regulation does not occur locally, but because such changes are insufficient to offset the perturbation. With local feedback, widespread compensatory changes develop initially but eventually reverse as the local input-output relationship is normalized (see below). If the input-output relationship of the perturbed compartment cannot be normalized, non-local compensatory changes persist. This is an important consideration for experimental manipulations, which are often very strong. By measuring compensatory changes after the perturbation is removed, one does not know if the compensation successfully offset the perturbation, but no compensatory change is likely to restore spiking in the presence of TTX.

Our results illustrate how the input-output relationships for different subcellular compartments are affected depending on whether compensation is directed by a local or global error signal. When regulation is local (i.e. compartment-specific), the effect of a perturbation is offset by compensatory changes in the affected compartment; compensatory changes start to develop in unperturbed compartments but reverse as output from the perturbed compartment is normalized through local compensation. In that scenario, the input-output relationship of the perturbed compartment is modified by persistent homeostatic changes whereas other compartments are ultimately unchanged. By comparison, if regulation is non-local, the global input-output relationship is returned to normal via compensatory changes distributed across compartments. In this scenario, the input-output relationship of the perturbed compartment is not returned to normal, and the input-output relationships in other compartments are adjusted away from normal. In short, restoring the global input-output relationship with non-local regulation distorts local input-output relationships, which could have important – likely negative – consequences for neural processing.

Regulating local input-output relationships is important if neurons operate as multicompartment processors rather than single computational units. Indeed, dendrites are increasingly recognized as semi-autonomous subunits capable of nonlinear integration and localized plasticity^41,42^. Our results suggest spatially restricted calcium transients may act as local error signals, enabling dendritic compartments to adjust synaptic weights and excitability independent of other compartments. Such modularity enhances computational flexibility, allowing localized plasticity without destabilizing global neuronal output^43^. From a control theory perspective, our simulations illustrate how local feedback loops confer robustness to perturbations. Prior modeling studies have shown that global feedback mechanisms can destabilize network activity unless finely tuned^44^. Our results suggest biological systems favor slow, compartmentalized homeostatic mechanisms to maintain stability across scales^45^. This principle parallels insights from network control theory, which emphasize the importance of localized feedback for sustaining dynamic stability in complex systems^46^.

Our results highlight the limitations of soma-centric feedback mechanisms, namely that they are liable to induce widespread compensatory changes that distort the local input-output relationships of functionally important subcellular compartments. For local compensation to occur, not only must error signals be local, those signals must also drive mechanisms that modulate ion channel expression (or function) locally. Evidence suggests that local protein translation, mediated by RNA-binding proteins such as FMRP and TDP-43, plays a critical role in homeostatic regulation^47,48^. Disruption of these localized processes has been implicated in disorders including Fragile X Syndrome and ALS, which hints at the potential importance of spatially precise homeostatic regulation^49^. Trafficking channels to or from the membrane is also likely be important^18^ and could conceivably be regulated locally^50^. Notably, some reports suggest transcriptional regulation as the primary driver^11^. Transcriptional changes seem more likely to mediate non-local compensation, although specificity could conceivably be achieved through a process like synaptic tagging^51^. Transcriptional, translational, and post-translational mechanisms are not mutually exclusive, and their intersection may be important, but their compartmental specificity still hinges on a local error signal.

In theory, intracellular calcium might simultaneously encode (multiplex) two error signals using fast and slow fluctuations and downstream effectors could extract (demultiplex) each signal by operating as high- or low-pass filters^12^. Indeed, theoretical work shows that combining a nonlinear intrinsic mechanism with a synaptic mechanism allows a single firing-rate-dependent signal to control both the mean and variance of firing, which is impossible for a single linear feedback^52–55^. Rather than extracting multiple features from a spectrally complex signal, spatially segregated calcium signals supply physically separate signals to separate effectors. Both strategies expand the effective degrees of freedom available for homeostatic control, and are not mutually exclusive. Other multiplexing strategies also conceivable.

Our simulation results suggest that calcium changes can be sufficiently segregated to support compartment-specific error signals. Experimental verification of this is difficult insofar as the calcium changes relevant for homeostatic regulation occur on long timescales and are thus difficult to stably track. Our results also suggest that less segregated calcium signaling can lead to increased cross-talk between feedback signals, which might contribute to pathology. For instance, loss of the A-type potassium channel gradient in dendrites, as reported in Fragile X Syndrome, enhances backpropagating action potentials and compromises calcium compartmentalization^33,56^. Similarly, impaired calcium buffering during aging and in Alzheimer’s disease has been associated with elevated cytosolic calcium, mitochondrial dysfunction, and synaptic degeneration^57,58^. These findings suggest that restoring calcium homeostasis and spatial segregation could offer therapeutic strategies to mitigate neuronal dysfunction that might, in part, arise from aberrant non-local homeostatic regulation.

Our simulations focussed on calcium-dependent feedback but neurons maintain homeostasis across multiple intracellular parameters, including pH, ATP/ADP ratio, and ionic composition^59,60^ that utilize feedback signals beyond calcium. Since many neurological disorders involve broad ionic dysregulation (Ca²⁺, Na⁺, K⁺, Cl⁻)^61,62^, incorporating these dimensions could refine models of homeostatic control under pathological conditions. Future modeling efforts should incorporate these additional dimensions to capture the full scope of homeostatic regulation. Mitochondrial health and the ATP/AMP ratio, for instance, are critical determinants of neuronal excitability and synaptic efficiency^63^. Integrating such metabolic and ionic feedback into multi-modal homeostatic models could yield a more comprehensive framework for understanding how neurons maintain stability across physiological and pathological contexts.

In summary, our simulations highlight the functional implications of implementing homeostatic changes in a compartment-specific manner and suggest that local homeostatic regulation is possible using spatially segregated activity-dependent calcium signaling. Our results also highlight the importance of certain methodological considerations, such as the perturbation strength.

## METHODS

### Non-spiking, two-compartment model

We started with an abstract non-spiking model comprising just two compartments (see Figure 1). Output (*o*) in the dendritic (*d*) or somatic (*s*) compartments was modeled as a function of input (*i*) using monotonic saturating functions in which parameters *α* and *β* control the “gain”:

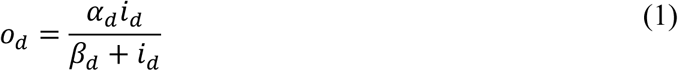

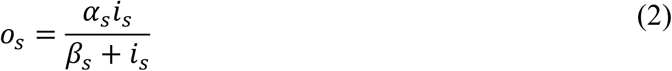

Parameters *α* and *β* are adjusted to homeostatically regulate the output of each compartment (see below). Dendritic output is scaled by a factor of *K_trans_* to produce the somatic input:

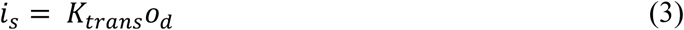

The difference between each compartment’s output and its target output constitutes an error signal, which dictates how gain control parameters are adjusted:

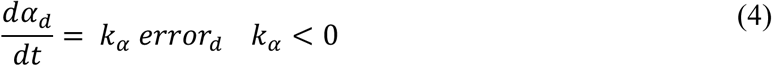

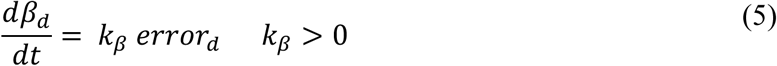

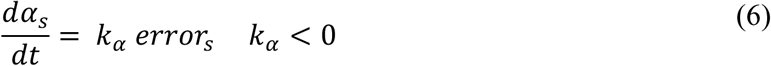

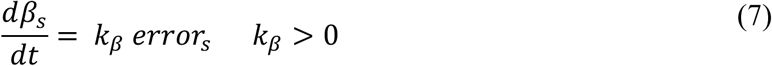

For the global feedback model, the *error* signal reflects both the somatic and dendritic errors:

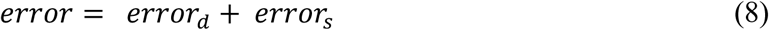

All parameter values for the non-spiking model are reported in Tables 1 and 2.

**Table 1.** – Parameter values for abstract 2-compartment model

| Parameter | Value |
| --- | --- |
| Dendritic compartment |  |
| $\alpha_d$ | 20 |
| $\beta_d$ | 100 |
| Activity-to-calcium time constant ($\tau_d$) | 0.1 |
| Baseline output ($o_{d,target}$) | 10 |
| Dendrite-to-soma attenuation factor ($K_{trans}$) | 20 |
| Somatic compartment |  |
| $\alpha_s$ | 20 |
| $\beta_s$ | 100 |
| Activity-to-calcium time constant ($\tau_s$) | 0.033 |
| Baseline output ($o_{s,target}$) | 1.82 |
| Feedback handling |  |
| Target output | 50 |
| $\alpha$ rate constant ($k_\alpha$) | 0.1 |
| $\beta$ rate constant ($k_\beta$) | -0.1 |

**Table 2.** – Values describing dendritic and somatic perturbations

| Perturbation | Parameter change |
| --- | --- |
| Increased dendritic | $i_d$: 10 → 50 |
| Increased somatic | $K_{trans}$: 20 → 50 |
| Decreased dendritic | $i_d$: 10 → 5 |
| Decreased somatic | $K_{trans}$: 20 → 10 |

**Table 3.**
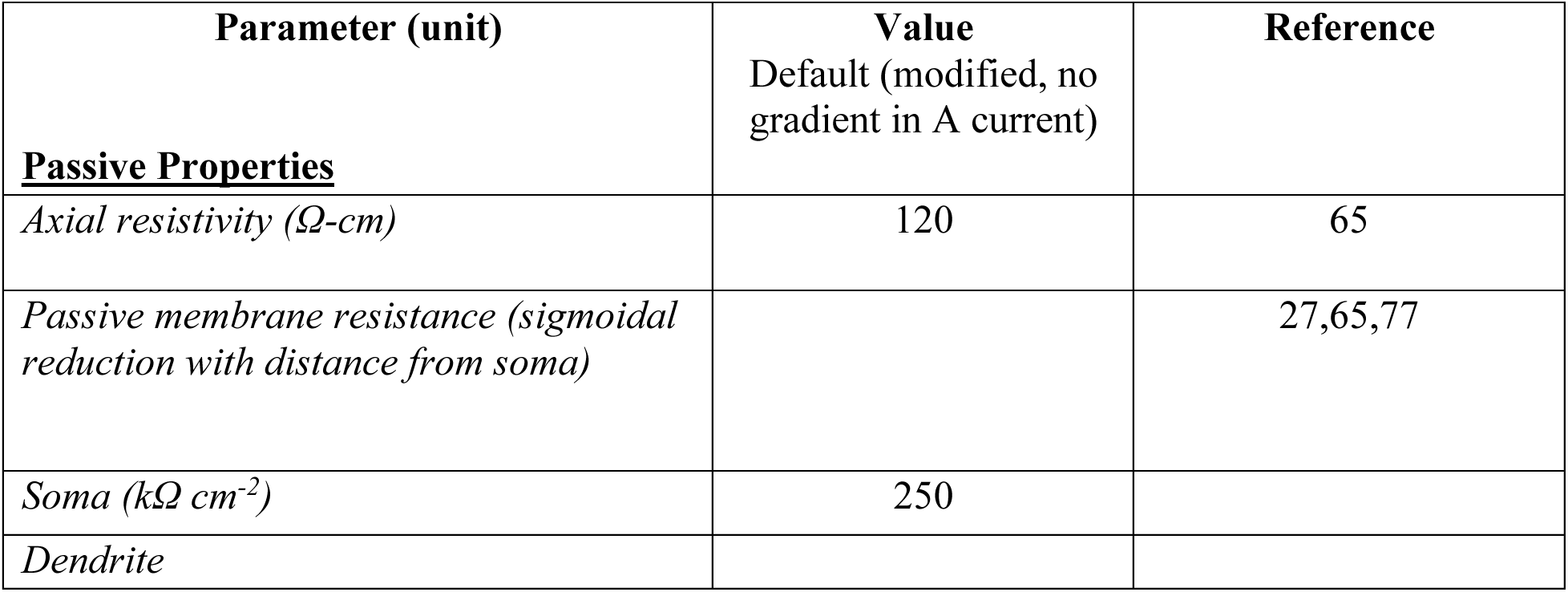
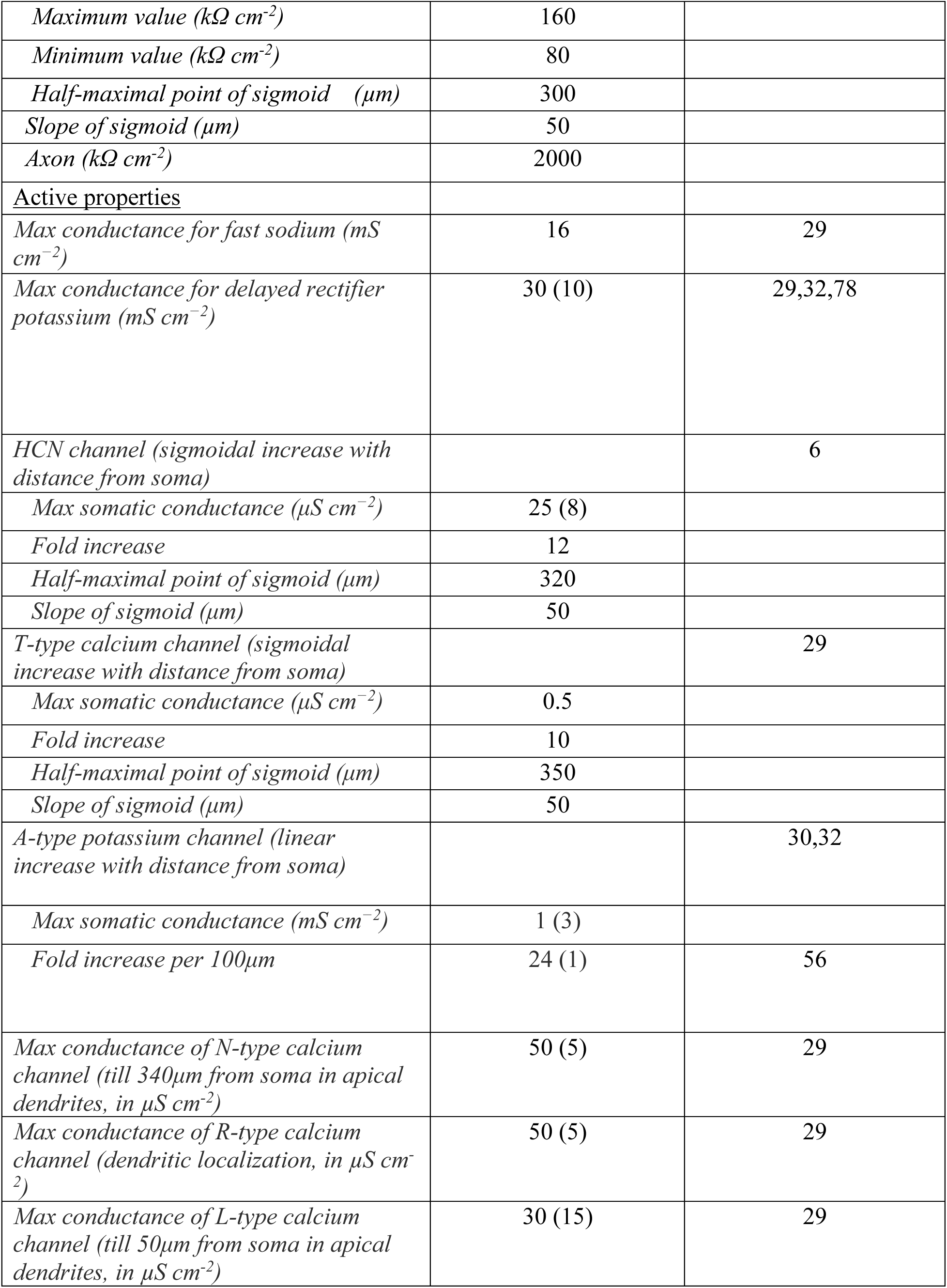
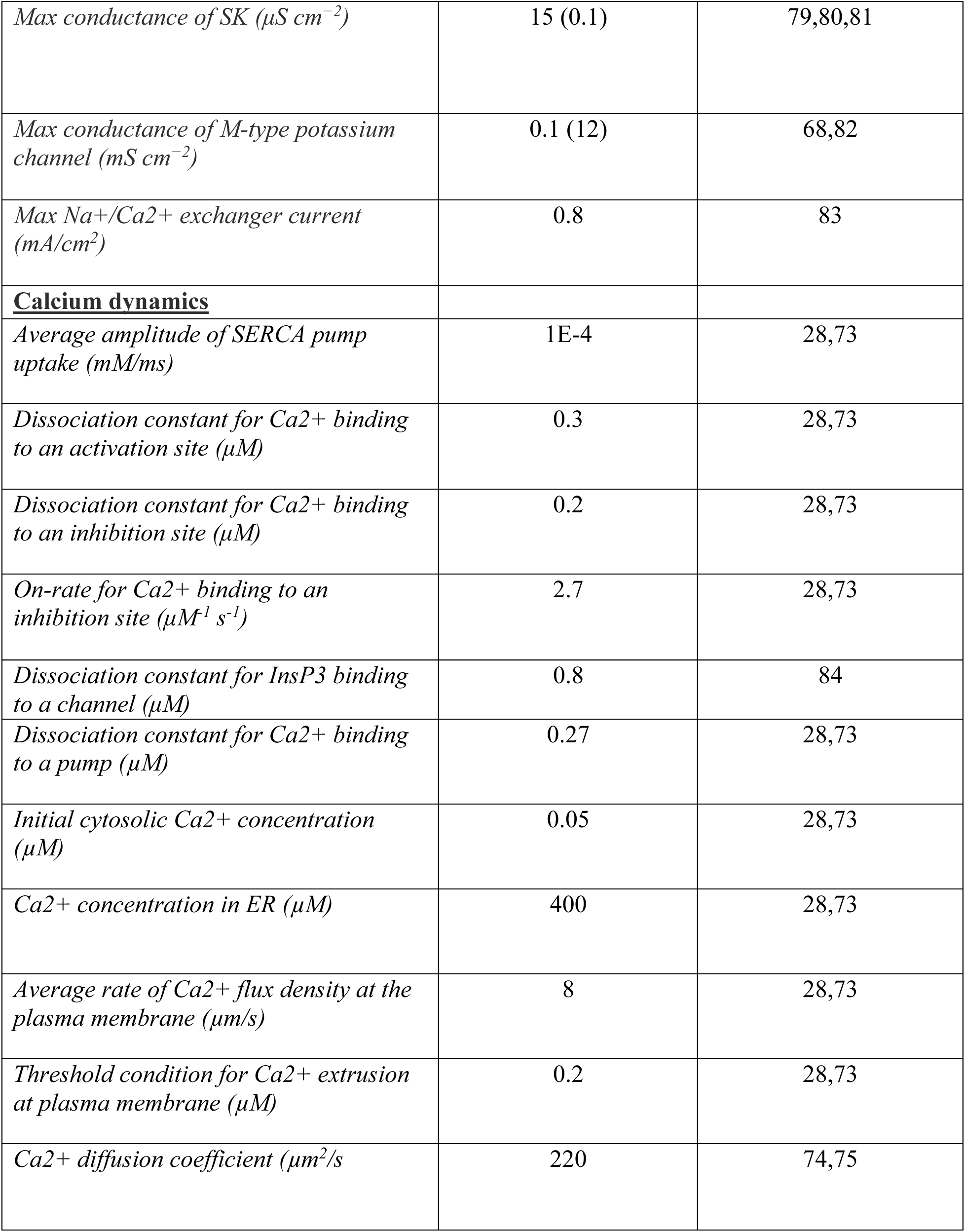
– Baseline parameter values for biologically realistic model neuron

| Parameter (unit) | Value<br>Default (modified, no<br>gradient in A current) | Reference |
| --- | --- | --- |
| **<u>Passive Properties</u>** |  |  |
| *Axial resistivity ($\Omega\text{-cm}$)* | 120 | 65 |
| *Passive membrane resistance (sigmoidal*<br>*reduction with distance from soma)* |  | 27,65,77 |
| *Soma ($k\Omega\text{ cm}^{-2}$)* | 250 |  |
| *Dendrite* |  |  |
| *Maximum value ($k\Omega\text{ cm}^{-2}$)* | 160 |  |
| *Minimum value ($k\Omega\text{ cm}^{-2}$)* | 80 |  |
| *Half-maximal point of sigmoid ($\mu\text{m}$)* | 300 |  |
| *Slope of sigmoid ($\mu\text{m}$)* | 50 |  |
| *Axon ($k\Omega\text{ cm}^{-2}$)* | 2000 |  |
| <u>Active properties</u> |  |  |
| *Max conductance for fast sodium ($\text{mS cm}^{-2}$)* | 16 | 29 |
| *Max conductance for delayed rectifier potassium ($\text{mS cm}^{-2}$)* | 30 (10) | 29,32,78 |
| *HCN channel (sigmoidal increase with distance from soma)* |  | 6 |
| *Max somatic conductance ($\mu\text{S cm}^{-2}$)* | 25 (8) |  |
| *Fold increase* | 12 |  |
| *Half-maximal point of sigmoid ($\mu\text{m}$)* | 320 |  |
| *Slope of sigmoid ($\mu\text{m}$)* | 50 |  |
| *T-type calcium channel (sigmoidal increase with distance from soma)* |  | 29 |
| *Max somatic conductance ($\mu\text{S cm}^{-2}$)* | 0.5 |  |
| *Fold increase* | 10 |  |
| *Half-maximal point of sigmoid ($\mu\text{m}$)* | 350 |  |
| *Slope of sigmoid ($\mu\text{m}$)* | 50 |  |
| *A-type potassium channel (linear increase with distance from soma)* |  | 30,32 |
| *Max somatic conductance ($\text{mS cm}^{-2}$)* | 1 (3) |  |
| *Fold increase per $100\mu\text{m}$* | 24 (1) | 56 |
| *Max conductance of N-type calcium channel (till $340\mu\text{m}$ from soma in apical dendrites, in $\mu\text{S cm}^{-2}$)* | 50 (5) | 29 |
| *Max conductance of R-type calcium channel (dendritic localization, in $\mu\text{S cm}^{-2}$)* | 50 (5) | 29 |
| *Max conductance of L-type calcium channel (till $50\mu\text{m}$ from soma in apical dendrites, in $\mu\text{S cm}^{-2}$)* | 30 (15) | 29 |
| *Max conductance of SK ($\mu S cm^{-2}$)* | 15 (0.1) | 79,80,81 |
| *Max conductance of M-type potassium channel ($mS cm^{-2}$)* | 0.1 (12) | 68,82 |
| *Max Na<sup>+</sup>/Ca<sup>2+</sup> exchanger current ($mA/cm^2$)* | 0.8 | 83 |
| **<u>Calcium dynamics</u>** |  |  |
| *Average amplitude of SERCA pump uptake ($mM/ms$)* | 1E-4 | 28,73 |
| *Dissociation constant for Ca<sup>2+</sup> binding to an activation site ($\mu M$)* | 0.3 | 28,73 |
| *Dissociation constant for Ca<sup>2+</sup> binding to an inhibition site ($\mu M$)* | 0.2 | 28,73 |
| *On-rate for Ca<sup>2+</sup> binding to an inhibition site ($\mu M^{-1} s^{-1}$)* | 2.7 | 28,73 |
| *Dissociation constant for InsP3 binding to a channel ($\mu M$)* | 0.8 | 84 |
| *Dissociation constant for Ca<sup>2+</sup> binding to a pump ($\mu M$)* | 0.27 | 28,73 |
| *Initial cytosolic Ca<sup>2+</sup> concentration ($\mu M$)* | 0.05 | 28,73 |
| *Ca<sup>2+</sup> concentration in ER ($\mu M$)* | 400 | 28,73 |
| *Average rate of Ca<sup>2+</sup> flux density at the plasma membrane ($\mu m/s$)* | 8 | 28,73 |
| *Threshold condition for Ca<sup>2+</sup> extrusion at plasma membrane ($\mu M$)* | 0.2 | 28,73 |
| *Ca<sup>2+</sup> diffusion coefficient ($\mu m^2/s$)* | 220 | 74,75 |

### Biophysically realistic multicompartment neuron model

We employed a morphologically realistic multicompartmental model of a CA1 pyramidal neuron with dendrites, soma, axon initial segment (AIS), and a simple axon. The morphology was modified from Neuromorpho.org^27,64^. Passive membrane properties arising from the cell membrane were represented as a capacitive current, and leak channels as a resistive current. The passive electrical behavior of the neuron was governed by three parameters: axial resistivity (*R_a_*), specific membrane resistivity (*R_m_*), and specific membrane capacitance (*C_m_*). In the base model, *R_a_* and *C_m_* were assigned a uniform value across the neuron (Table 3) whereas *R_m_* was spatially nonuniform, varying as a sigmoidal function of distance from the soma (*x*), consistent with Basak and Narayanan^65^,

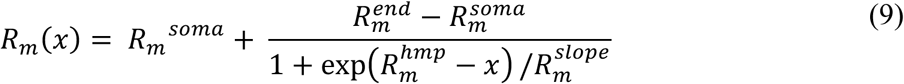

The neuron model was segmented using the *d_λ_* rule^66^ such that each segment was shorter than one-tenth of *λ*, the space constant at 100 Hz; in the base model, this resulted in 897 distinct segments.

We incorporated voltage-gated sodium, potassium, and calcium channels with gating properties taken from previous electrophysiological studies (see Table 3). The model also includes calcium-gated small conductance potassium (SK) channels, Na^+^/Ca^2+^ exchanger, IP_3_-mediated intracellular calcium release, calcium pumps, calcium buffers with their spatial distribution matching experimental data (see Table 3). Reversal potentials for *Na*^1^, *K*^1^, and ℎ channels were 55, −90, and −30 mV, respectively. Calcium currents, by contrast, were modeled using the Goldman–Hodgkin–Katz (GHK) formalism, with intracellular and extracellular calcium concentrations of 50 nM and 2 mM, respectively.

These channels were distributed along the somatodendritic axis to match experimental recordings (see Table 3 and Fig. 5A). Specifically, fast sodium and delayed-rectifier potassium channels were distributed uniformly. The *A*-type potassium channel density increased linearly as a function of distance from the soma:

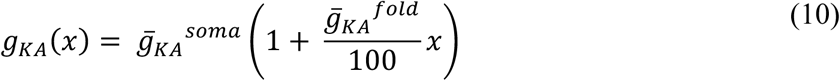

The HCN and *T*-type calcium channel densities were modeled as sigmoidal functions, increasing with radial distance from the soma:

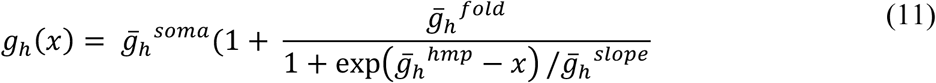

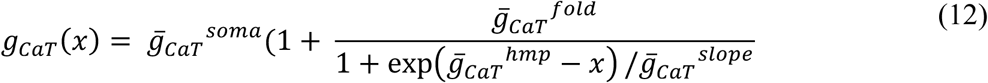

The *M*-type potassium and *L*-type calcium channels were perisomatic^29,69^. The SK and the *R*-type calcium channels were distributed uniformly across the apical dendrites. The *N*-type calcium channels were distributed uniformly up to 340 µM radial distance along the apical dendrite. Distances are specified as radial distances from the soma to match experimental measurements, which are conventionally reported as radial rather than path distances.

The model contains 150 colocalized AMPAR and NMDAR synapses distributed randomly over the apical dendrites of the neuron with NMDAR-to-AMPAR ratio of 1:1^70^. The current through the NMDAR was partitioned into components carried by three ionic species: Na, K, and Ca^2+^. The voltage- and time-dependence of each component was modeled using the GHK formulation^28,65,71,72^:

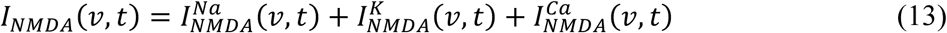

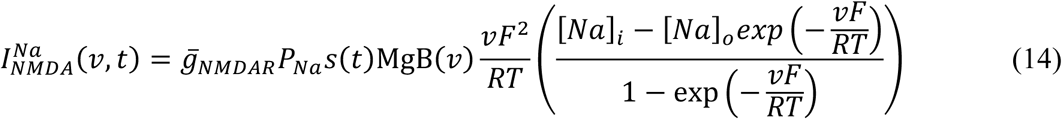

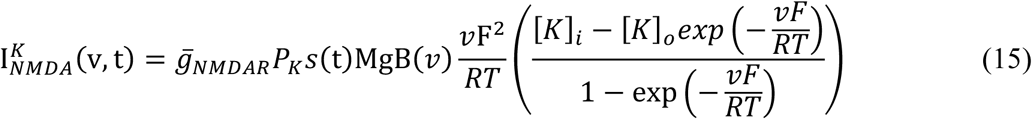

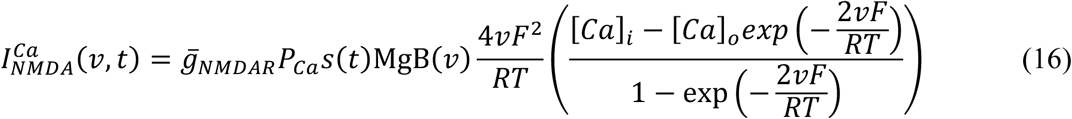

Here, *g̅_N_*_MDA<_defined the maximum conductance of NMDA receptors. The relative permeability ratios were set to *P_Ca_* = 10.6, *P_Na_* = 1 and *P*_K_ = 1. The ionic concentrations were set to [*Na*]*_i_* = 18 mM, [*Na*]*_o_* = 140 mM, [*K*]*_i_* = 140 mM, [*K*]*_o_* = 5 mM, [*Ca*]*_i_* = 100 nM and [*Ca*]*_o_* = 2 mM. The magnesium dependence of the NMDAR current:

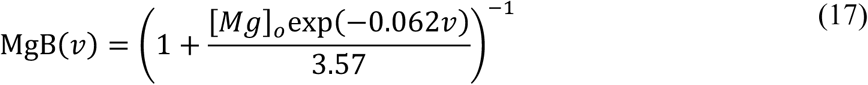

with [*Mg*]*_o_* = 2 mM. The kinetics of the NMDAR current were governed by *s*(*t*):

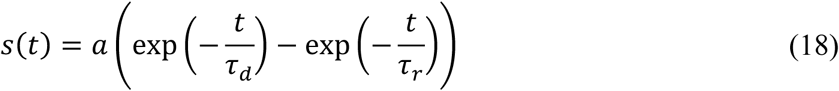

Here, *a* is a normalization constant ensuring 0 ≤ *s*(*t*) ≤ 1, the decay time constant *τ_d_* = 50 ms, and the rise time *τ_r_* = 5 ms.

The current through the AMPA receptor was mediated by two ions, Na^+^ and K^+^:

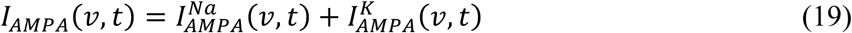

with the sodium and potassium components given by:

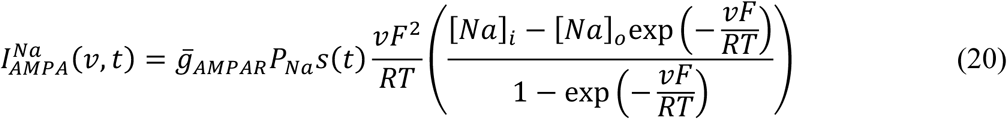

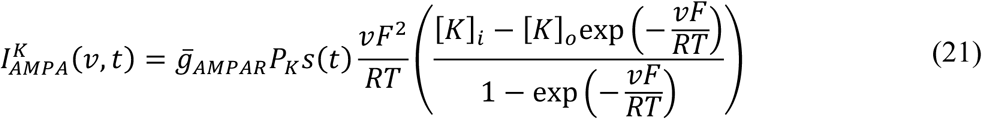

The relative permeability ratios were set to *P_Na_* = 1 and *P_K_* = 1, where *g̅_AMPA_*_<_defines the maximum conductance of AMPA receptors. *s*(*t*) was modeled as for the NMDAR (Eqn 18) but with τ*_r_* = 2 ms and τ*_d_* = 10 ms. To normalize the unitary EPSP associated with each synapse, we ensured that attenuation along the dendritic cable did not affect the unitary somatic EPSP amplitude. Accordingly, *g̅_AMPA_*_<_along the somatoapical trunk was tuned to produce a unitary somatic response of ∼0.2 mV irrespective of the synaptic location^31^.

Parameter values (see Table 3) were tuned to match several physiological response properties including input resistance and firing rate^6,27^. Spontaneous firing of 1-2 Hz was produced by stimulating each synapse with independent Poisson processes at 8 Hz.

### Calcium dynamics

Overall Ca^2+^ dynamics were modeled as the combined contribution of the various mechanisms affecting cytosolic Ca^2+^ concentration ([Ca^2+^]_c_). The partial differential equation governing cytosolic Ca^2+^ dynamics^73^:

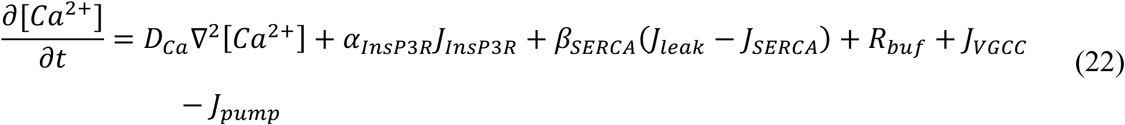

where *D*_Ca_ is the experimentally determined diffusion coefficient for calcium^74,75^, *α_InsP_*_H*R*_ is the density gradient of InsP_3_Rs along the ER, and *β_SERCA_* is the density of SERCA pumps and leak channels on the ER along the somatodendritic axis. *J*_InsP3R_, *J*_VGCC_, *J*_SERCA_, *R*_buf_, *J*_pump_, and *J*_leak_ denote the Ca^2+^ fluxes due to InsP3Rs, VGCCs, SERCA pumps, buffering, plasma-membrane Ca^2+^ extrusion pumps, and ER leak channels, respectively. Radial Ca^2+^ diffusion was implemented by dividing each neuronal compartment into four concentric annuli, and diffusion along the longitudinal axis of the neuron was also included^66^. Each flux is modeled like in Ashhad and Narayanan^28^.

A calcium-dependent feedback process adjusts ion channel densities according to Eqn 23:

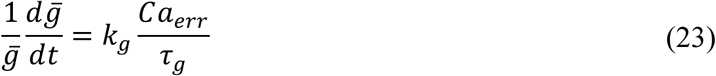

where *g̅* is channel density and *k*_K_ is a scaling factor (see below for which channels are adjusted). The calcium-based error signal (*Ca_err_*) reflects deviation of calcium from its target value:

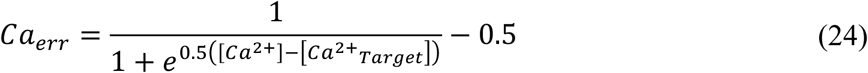

where *τ*_K_ is the integration time constant. Our approach addresses an inherent asymmetry in the calcium-based error signal and, consequently, in the homeostatic compensation. For a typical set point of 70 nM intracellular free calcium, deviations below the set point are mechanistically bounded at approximately 50 nM, whereas deviations above it can extend into the micromolar range. In the absence of a saturating function, this asymmetry would produce disproportionately large compensatory changes in ion channel densities in response to calcium influx. Incorporating a sigmoidal function constrains the error signal at the extremes and confers approximately equal weighting to increases and decreases in calcium concentration, thus promoting stable homeostatic regulation.

Average calcium concentration can differ across compartments. *Ca_err_* was calculated with 70 nM as target calcium but, for the soma, *Ca_err_* was set to zero if [*Ca*^21^] was between 65 and 75 nM, thus encouraging stability of the model. The range was expanded to 65 – 85 nM for dendrites. These ranges were decided based on the calcium rise due to local perturbations. Feedback was applied to AMPAR and NMDAR to implement synaptic scaling, and to the fast sodium channel and M-type potassium channel to implement excitability regulation. Simulations always started with parameters at baseline values (see Table 3) after which only values subject to feedback were adjusted. The *k*_K_ values were chosen such that when fast sodium channel or AMPAR/NMDAR density increases, M-current density decreases, and vice versa (see Table 4). The regulation time constants were set to 40 or 50 s (see Table 4), which is much slower than individual spikes or synaptic events, but is still much faster than the hours over which homeostatic regulation normally occurs. This was for practical reasons, to reduce simulation time.

**Table 4.**
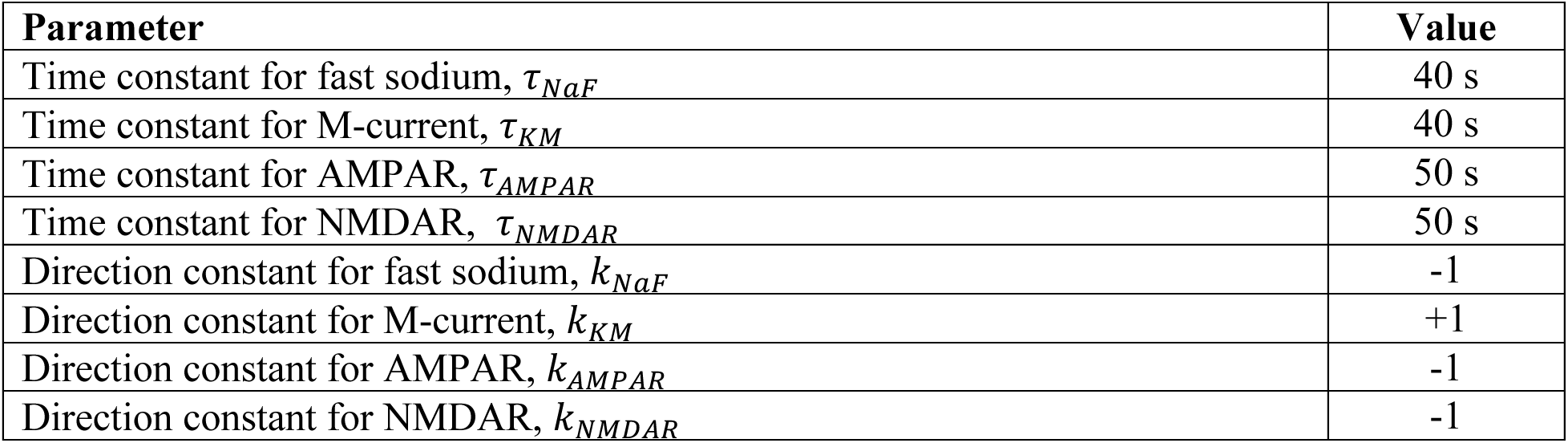
– Calcium-dependent regulation parameters

| **Parameter** | **Value** |
| --- | --- |
| Time constant for fast sodium, $\tau_{NaF}$ | 40 s |
| Time constant for M-current, $\tau_{KM}$ | 40 s |
| Time constant for AMPAR, $\tau_{AMPA}$ | 50 s |
| Time constant for NMDAR, $\tau_{NMDAR}$ | 50 s |
| Direction constant for fast sodium, $k_{NaF}$ | -1 |
| Direction constant for M-current, $k_{KM}$ | +1 |
| Direction constant for AMPAR, $k_{AMPA}$ | -1 |
| Direction constant for NMDAR, $k_{NMDAR}$ | -1 |

The neuron was subjected to four different perturbations: (*i*) Increased rate of synaptic input to dendrites, where 50 of the 150 synapses (in three oblique dendrites; the red dendrite marked in **Fig 5A** is one of the three stimulated dendrites) were stimulated at a higher frequency of 20 Hz. Input was increased across three dendrites since adjusting input to a single dendrite could not cause a notable increase in firing rate. (*ii*) Decreased synaptic strength, where AMPAR and NMDAR weights were reduced by 50%. The change in *ii* is unlike that in *i* since reducing synaptic input to zero in the three dendrites perturbed in *i* did not cause a notable change in the somatic firing; thus, to simulate a low synaptic input scenario in *ii*, synaptic input across all synapses was reduced by 50%. (*iii*) Increased somatic excitability, where the soma was stimulated with 20 pA along with background synaptic input. (*iv*) Decreased somatic excitability where a leak conductance with -70 mV as reversal potential (*E_rev_*) was introduced into the soma (*g̅_in_*_ℎ_= 0.1 mS/cm^2^) to simulate increased perisomatic inhibition:

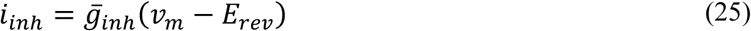

For all perturbations, the membrane potential, internal calcium concentration, and channel densities were recorded from the soma, one of the perturbed dendrites and one nearby oblique dendrite that was not perturbed.

### Modified model with uniformly distributed in A-type potassium channels

In the soma, most calcium influx is due to spiking whereas most of the calcium influx in dendrites is because of synaptic input involving NMDARs^24,76^. Back propagating action potentials (bAPs) also contribute to calcium influx in dendrites by activating high voltage-activated calcium (HVACa) channels. Due to the spatial distribution of A-type potassium channels in the dendrites, bAPs attenuate in amplitude as they travel across the dendrite to distal locations^30,32^, which mitigates the activation of HVACa channels, rendering synaptic activity the major contributor of dendritic calcium. To reduce the extent of spatial segregation of calcium inside the neuron for some simulations (see Fig. 8), we replaced the gradient of A-type potassium channel density with a uniform density. This led to reduced attenuation of bAP amplitude, thereby increasing spike-dependent calcium influx in distal dendrites. This change also increases overall excitability, which prompted us to revise parameters (see Table 3) to match the initial baseline spontaneous firing rate but without changing the new bAP attenuation pattern. Reduction of dendritic A-type potassium channels also led to a higher amplitude of dendritic spikes, which travel to the soma, enabling distal synaptic input to have a greater impact on somatic calcium. This led to an increase in correlation of somatic and dendritic calcium or, in other words, reduced spatial segregation of calcium.

### Code/Software

Simulations for the non-spiking abstract model were conducted in Matlab (R2025b), while morphologically realistic models were run using the NEURON environment (Version 8). All code will be publicly accessible on GitHub after publication.

## Contributions

AR and SAP conceived the study. AR conducted all simulations and analysis. AR and SAP prepared the manuscript.

## Acknowledgements

This study was funded by a Discovery Grant (RGPIN 2024-05431) from the Natural Sciences and Engineering Research Council of Canada to SAP. AR was also supported by the Yuet Ngor Wong Fellowship from the University of Toronto.

---

**Figure S1.**
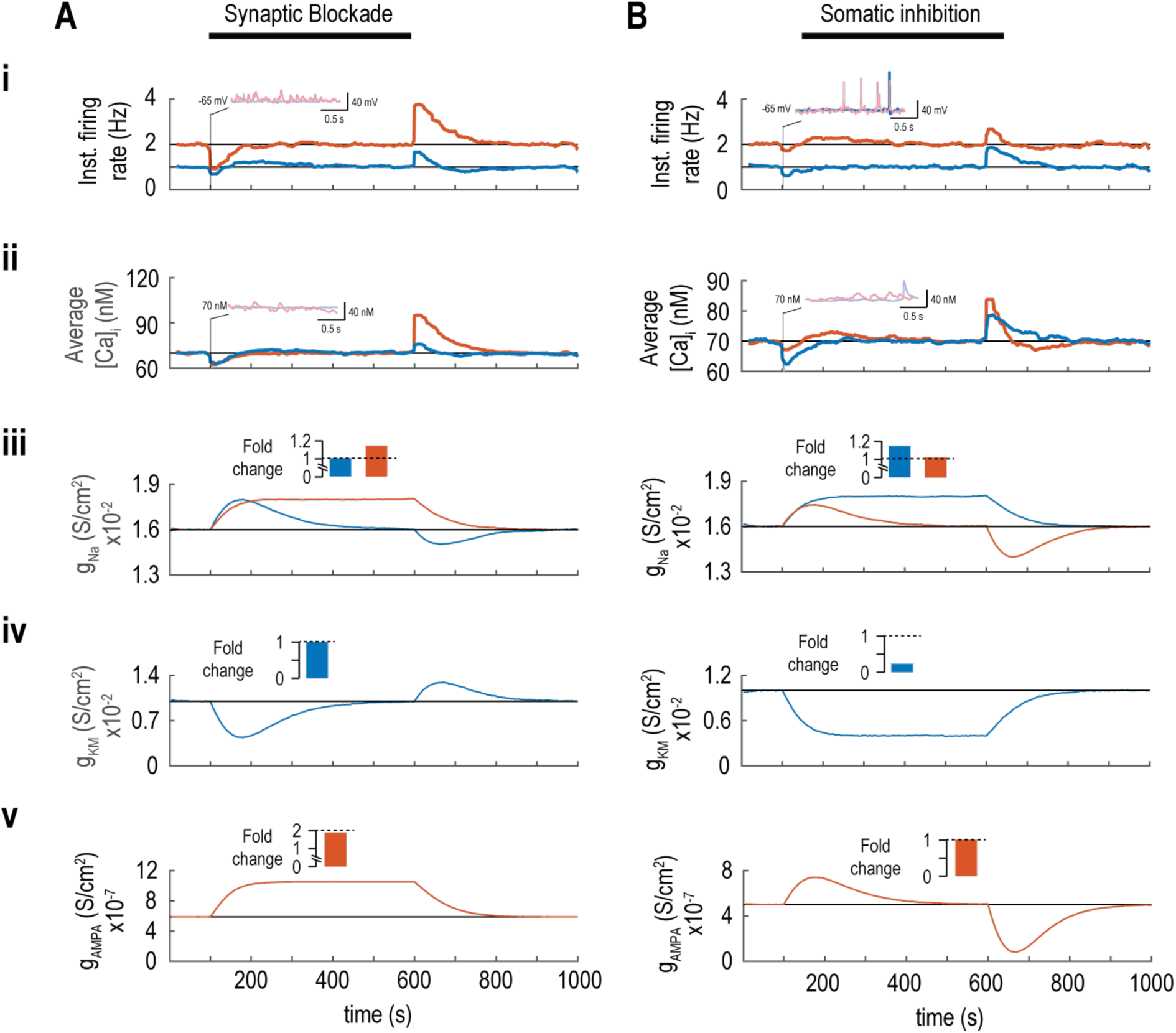
– Firing rate, calcium and conductance density for decreased input to dendrite and decreased activity in soma for realistic spiking model. Compare to increased input in Figure 6. Unlike in Figures 6-9, yellow traces are not shown here since the synaptic perturbation was applied to all dendrites, meaning the yellow trace no longer constitutes as unperturbed control for comparison with the perturbed dendrite (red); instead, both dendritic locations exhibited similar changes, consistent with the spatial extent of the perturbation. **A.** Synaptic blockade. Firing rate (i) and intracellular calcium (ii) were reduced by the perturbation but returned to baseline following compensatory changes. Compensatory changes in *g̅*_Na_ (iii) and *g̅*_AMPA_ (v) were sustained in the dendrites whereas the soma exhibited only a transient change in *g̅*_Na_(iii) and *g̅*_KM_(iv). The fold change in conductance densities at steady state (i.e. just prior to termination of the perturbation) are summarized as bar graph insets. **B.** Somatic inhibition. Data, plotted like in A, reveal that the firing rate and calcium in each compartment were returned to normal by sustained compensatory changes in *g̅*_Na_, *g̅*_KM_ and *g̅*_AMPA_ in all compartments.

**Figure S2.**
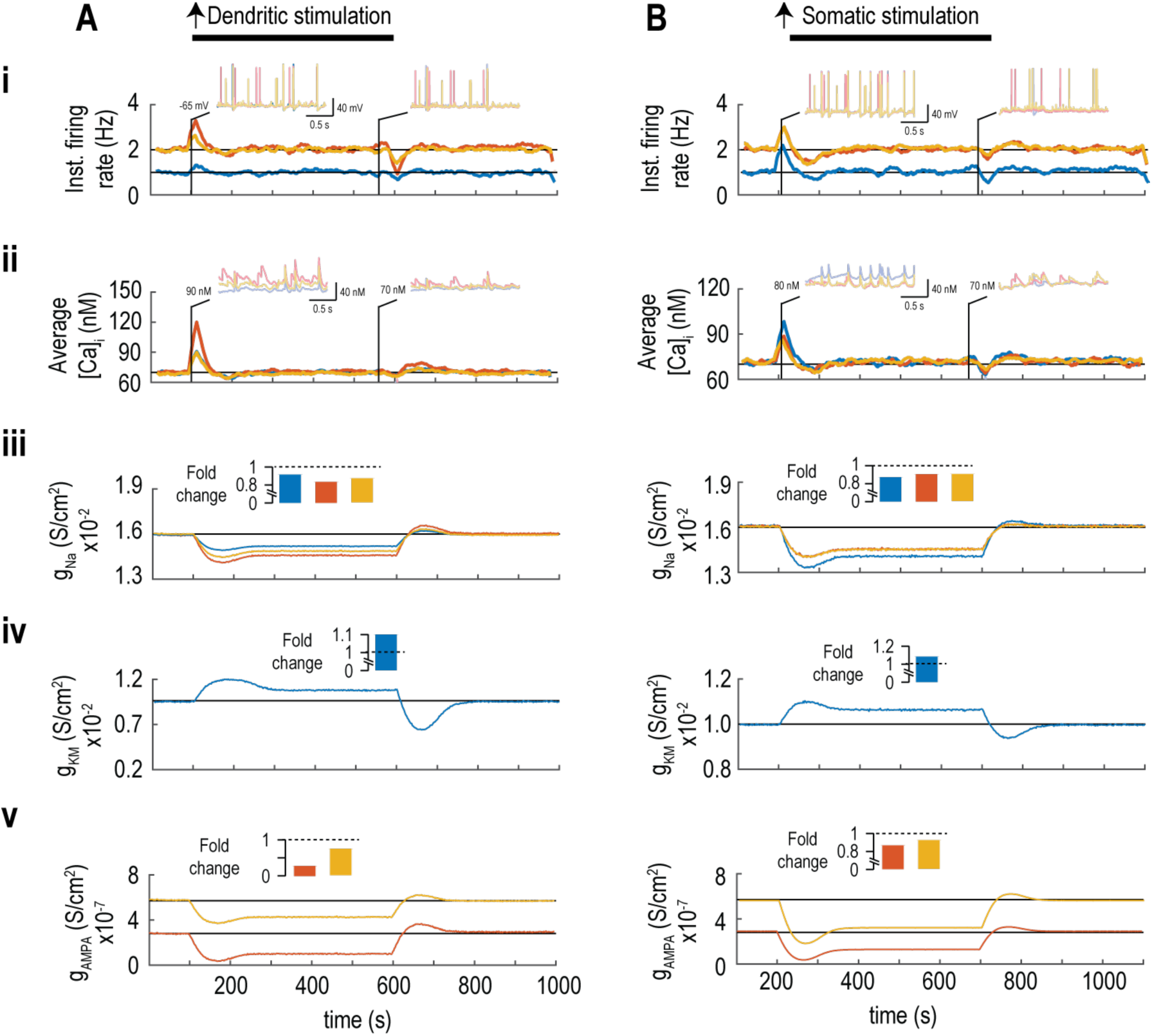
– Firing rate, calcium and conductance density for increased input to soma and dendrite for real spiking model with no gradient in *g*s_KA_. Compare to reduced input in Figure 8. **A.** Increased dendritic stimulation. Firing rate (i) and intracellular calcium (ii) were altered by the perturbation but both returned to baseline following compensatory changes. Compensatory changes in *g̅*_Na_(iii), *g̅*_KM_ (iv), and *g̅*_AMPA_ (v) were sustained in all compartments, reflecting non-local homeostatic changes due to compromised calcium segregation in this model. The fold change in conductance densities at steady state (i.e. just prior to termination of the perturbation) are summarized as bar graph insets. **B.** Increased somatic stimulation. Data, plotted like in A, reveal that the firing rate and calcium in each compartment were returned to baseline by sustained compensatory changes in *g̅*_Na_, *g̅*_KM_ and *g̅*_AMPA_ in all compartments, again reflecting non-local homeostatic changes.

